# Comprehensive analysis of ammonia and biogenic amines in ecologically diverse systems with high-resolution mass spectrometry

**DOI:** 10.64898/2026.09.22.753493

**Authors:** Mateusz Fido, Elisa Cappio Barazzone, Etienne Hoesli, Jens Scheidegger, Nimisha Khurana, Jeongmin Kim, Markus Arnoldini, Martin Sperfeld, Raffael Luca Schumann, Shinichi Sunagawa, Annika Hausmann, Renato Zenobi, Emma Slack

## Abstract

The acquisition of sufficient nitrogen for growth, as well as the disposal of highly toxic nitrogen-containing degradation products, represent serious metabolic challenges in all multicellular organisms and microbial ecosystems. Understanding these processes requires a broadly applicable system to accurately quantify the concentration of a wide spectrum of biological amines and inorganic nitrogen-containing compounds. To date, this has been challenging due to limited specificity, selectivity, breadth and/or sensitivity of assays. Here we present a simple, optimized method based on liquid chromatography and high-resolution mass spectrometry, with an accompanying analysis pipeline, for the accurate quantification of more than 30 of these compounds, including ammonia. We demonstrate the effectiveness of this method in quantifying relevant nitrogen-containing compounds in mouse and human gut content, bacterial cultures, algal symbiosis in a cnidarian and freshwater ecosystem samples. As a proof-of-concept, we applied the method to study the influence of the gut microbiome on cecal nitrogen distribution in mice with different gnotobiotic microbiomes. This revealed a profound influence of gut microbiota composition on amino acid, ammonia, and amine concentrations in the upper large intestine. Overall, we provide a broadly applicable approach that can be used across microbial ecology fields to generate novel insights into the biology of nitrogen assimilation, exchange, and disposal.

## 1 Introduction

Nitrogen-containing compounds are essential for life and play major roles in a wide range of metabolic and environmental biological processes. For example, ammonia is required for the formation of amino acids and nucleic acids, and is important in regulating the acid-base homeostasis in microbial systems.^1–3^ While animals also need to form amino acids and nucleic acids, elevated systemic levels of ammonia, referred to as hyperammonemia, are highly toxic, and can be lethal.^1,4,5^ Biogenic amines, a diverse group of compounds which include catecholamines, polyamines (agmatine, cadaverine, putrescine, spermine and spermidine), serotonin, tryptamine, and histamine, are present across different tissues and kingdoms of life, with functions ranging from neurotransmission to genome protection.^6–9^ Nitrogen exchange has also been suggested to underlie control of some symbioses, and disruption to this exchange can be detrimental to the host, for example coral bleaching driven by temperature stress.^10^ Consequently, a method for precise and accurate monitoring and quantification of these compounds across diverse matrices and concentration ranges would be highly relevant for diagnosing diseases, understanding metabolic states and disorders, and monitoring, understanding and preserving healthy agriculture and aquatic systems.^11,12^

To date, the available methods are complex or specific for a single application or compound, such as clinical blood ammonia measurements.^13–17^ Traditional techniques are based on colorimetric or fluorometric assays, which typically achieve detection limits in the micromolar range but often fail to differentiate between amine-containing compounds present in complex matrices.^18–20^ Gas chromatography (GC) coupled with mass spectrometry (MS) is a highly sensitive technique for detecting volatile nitrogen-containing compounds, with reported methods for the quantification of several biogenic amines^21^ and ammonium nitrate.^22^ Although GC-MS can achieve very low (ppb levels) limits of detection and quantification, it requires extensive sample preparation and cannot be applied directly to liquid-phase samples, which can constrain its use outside specialized analytical settings. Another common approach is ultra- and high-performance liquid chromatography (U/HPLC) coupled with absorbance, evaporative light scattering or fluorescence detection. This method can typically achieve micromolar detection limits but again suffers from selectivity issues in complex sample matrices containing interferants and isomers with similar chemical properties.^23^ More recently, electrochemical and optical biosensors have been developed for real-time monitoring of ammonia, offering even lower limits of detection; however, extensive development is required before such sensors become available and validated for a broad range of amine containing compounds.^12,24,25^

Liquid chromatography coupled to high-resolution mass spectrometry (LC-HRMS) combines the advantages of improved chromatographic separation, high specificity and sensitivity, with the unparalleled selectivity and quantifiability of high-resolution mass spectrometry. Numerous studies have explored derivatization strategies for nitrogen-containing compounds.^22,26–32^ One common reagent, used primarily for the derivatization of amino acids, is diethyl ethoxymethylene-malonate (DEEMM). DEEMM undergoes a chemoselective esterification reaction with primary and secondary amines (e.g., proline), resulting in the formation of stable aminoenone derivatives that absorb in the ultraviolet range.^33^ This reaction is catalyzed by a base, such as potassium hydroxide, and occurs under mild conditions (room temperature in water/methanol solutions). While initially developed for *N*-protection during the synthesis of amino sugars and amino acid esters,^30,34^ DEEMM was later adapted for detection of amines.^28,29,35–44^ Notably, the reaction does not exhibit unspecific byproducts.^41^ The main products are generally chemically stable over days to weeks, with the exception of the secondary amine derivatives, which have been shown to degrade significantly within 7 days.^33^

In this work, we demonstrate that DEEMM derivatization combined with LC-HRMS and appropriate data processing permits precise, sensitive and versatile quantification of amino acids, biogenic amines and ammonia in environmentally diverse samples. We optimized the technique for use in murine gastrointestinal tract content, but demonstrate its applicability to human fecal samples, spent bacterial media of *E. coli* fed different nitrogen sources, sea anemones and their algal symbionts, and environmental lotic and lentic fresh-water. We were able to accurately identify fifty biologically relevant compounds and reliably quantify more than thirty. Most notably, this included ammonia, amino acids, polyamines, neurotransmitters, nucleobases, catecholamines, and urea cycle intermediates. We found DEEMM derivatization efficiency to vary significantly with respect to the compound and sample matrix and devised a dedicated quantification pipeline implemented via an open-source, cross-platform mass spectrometry software, LCMSpector, to allow efficient targeted quantification of complex spectra.^45^ Compared to colorimetric assays, which can both overestimate or underestimate ammonia content when used for gut content, this approach reproducibly quantifies ammonia in this extremely complex matrix. We further demonstrate how data-dependent acquisition allows for tracing of DEEMM derivatives without losing information on the rest of the sample content. Finally, as a proof-of-concept we use the approach to determine how microbiome composition alters the concentration of nitrogen-containing compounds in the mouse cecum, revealing profound effects of the microbiome on a wide range of amines and ammonia.

## 2 Results

### 2.1 UPLC-UV-HRMS analysis of nitrogen-containing compounds

To evaluate the quantitative performance of DEEMM derivatization for our target compounds, we prepared chemically pure standards for 50 biologically relevant nitrogen-containing compounds (Supporting Information, Table S1) across a range of concentrations from 10 mM down to 1 µM. After derivatization, compounds were separated using reversed-phase chromatography on a standard C18 column and analyzed using an Orbitrap high-resolution mass spectrometer. Owing to the characteristic neutral loss fingerprint of DEEMM aminoenones, the annotation of singly substituted derivatives is straightforward. For most target compounds, the principal pseudomolecular ion [M+H]^+^ and the neutral loss pseudomolecular ion [M-C_2_H_6_O+H]^+^ showed comparable MS intensities. However, because DEEMM derivatization is not regioselective, all accessible nucleophilic amine residues can react with the derivatizing agent, producing higher-order derivatives, which must be accounted for during quantification (Supporting Information, Table S1). Considering the sterically accessible primary and secondary amine residues in each targeted compound, we compiled ion lists covering all the possible reaction products and their corresponding neutral loss ions (Supporting Information, Table S1). We then employed the in-house developed software LCMSpector to verify the selected ion chromatograms for each metabolite and generate linear regression curves for calibration.^45^ For all of the targeted compounds where sufficient quantitative signal was obtained, we adjusted the range to obtain linear detection curves over the full range of tested concentrations. More than 30 target compounds met the calibration linearity criterion of R^2^ ≥ 0.90 used for quantification.

Double-substituted derivatization products constitute on average more than 25% of total ion intensity of all traced compounds, and failure to account for them would lead to significant errors in quantitation. Considerable inter-sample variability was also observed in the relative ion composition of individual compounds, indicating that all potential ions should be included in analyses.

To further validate the accuracy of our LC-HRMS method, we benchmarked it against commonly used colorimetric and fluorimetric assays specifically for ammonia quantification (Fig. 2). We compared both commercial and in-house kits based on (i) the Berthelot (indophenol) reaction, which relies on the oxidative coupling of ammonia with phenolic compounds in the presence of hypochlorite, and (ii) the o-phthalaldehyde (OPA) reaction, in which ammonia reacts with OPA and a thiol co-reactant to form a fluorescent isoindole derivative.^46,47^ Although these assays are convenient and inexpensive, they showed poor linearity and inconsistent recovery across sample matrices (Supporting information - Fig. 1A), and either under- or over-estimated ammonia levels when compared to LC-MS analysis of the same samples (Fig. 2). Spike-in experiments using known amounts of ammonium chloride in plasma revealed apparent ammonia concentrations exceeding the expected values (Supporting information - Fig. 1B). This systematic overestimation is consistent with off-target reactivity of the colorimetric and fluorimetric reagents toward other primary amines, peptides, or chromogenic components abundant in biological matrices. Additional controls comparing protein precipitation and filtration failed to reduce the apparent signal, indicating that the background arises predominantly from small-molecule amines rather than proteins (Supporting information - Fig. S1C). By contrast, quantification by LC–HRMS showed agreement with the biologically expected concentrations^48^ and remained unaffected by sample matrix composition, confirming the specificity and robustness of this approach for measuring ammonia and other nitrogenous compounds.

**Figure 1.**
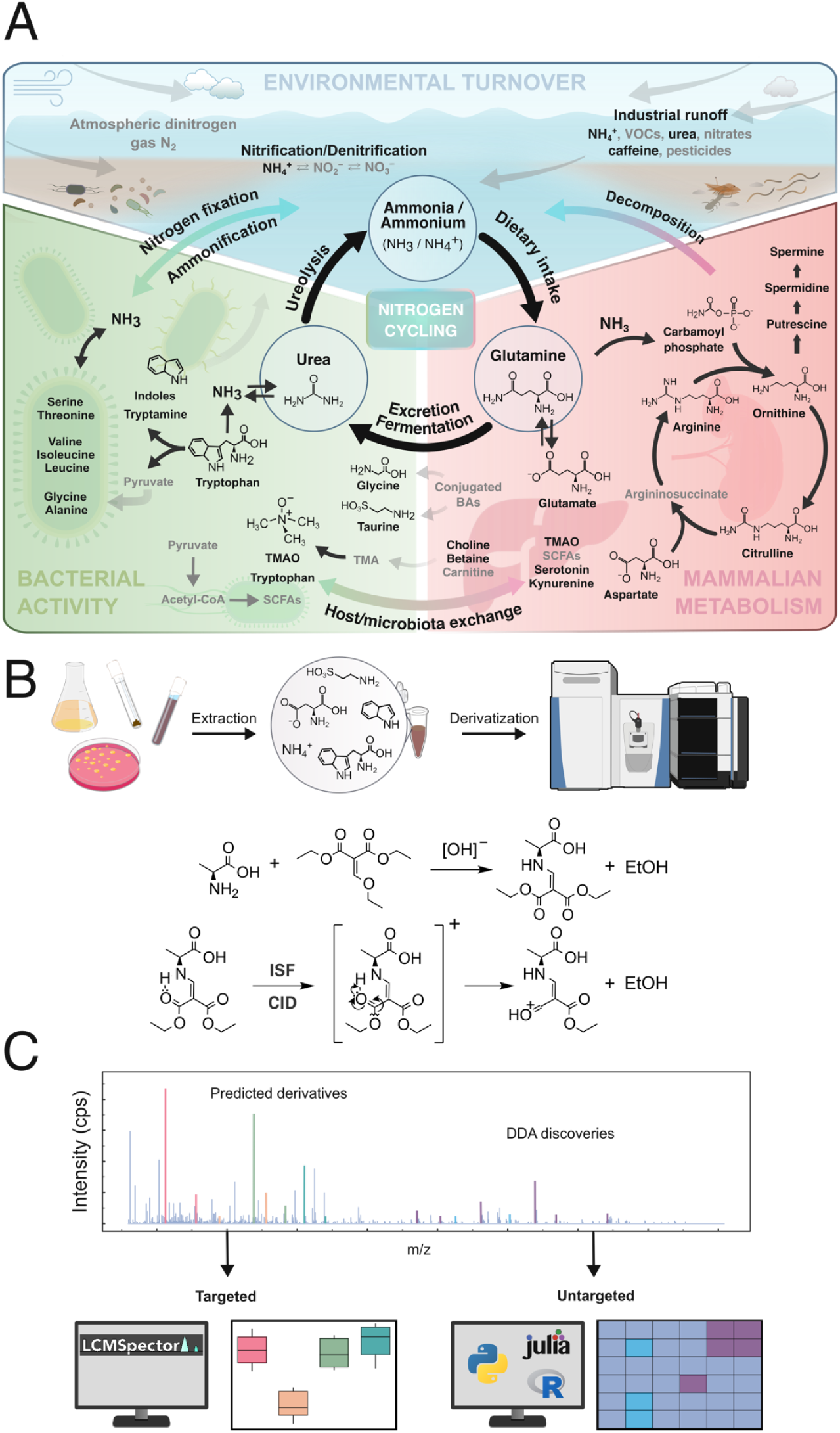
Nitrogen-containing compounds measurable using our LC-HRMS method. **A)** Targeted N-compounds (black) visualized on a simplified diagram of nitrogen cycling between mammalian metabolism, bacterial activity and environmental turnover. **B)** Nitrogen-containing compounds are extracted depending on the sample phase, derivatized with DEEMM, and measured via LC-HRMS using data-dependent acquisition (DDA). This allows for direct quantification of the predicted derivatives, as well as discovery and semi-quantitative workflows for untargeted molecules. **C)** Data-dependent acquisition allows for non-discriminatory identification of precursor ions present in the sample, enabling quantification of the targeted N-compounds, as well as discovery-based searches. Abbreviations: ISF – in-source fragmentation, CID – collision-induced dissociation, DDA – data-dependent acquisition, TMAO – trimethylamine oxide, TMA – trimethylamine, BA – bile acids, SCFA – short-chain fatty acids, VOC – volatile organic compound.

**Figure 2.**
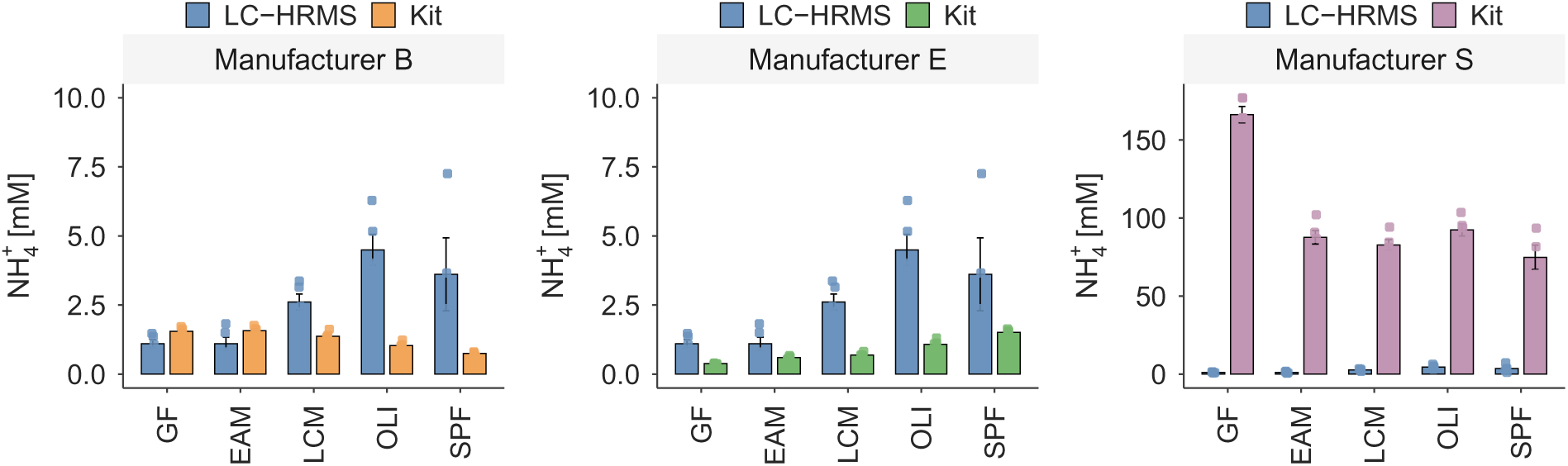
Comparative quantification of cecal ammonia concentrations using LC-HRMS and three commercial assay kits. Clustered bar plots show mean ammonia concentrations ± SEM measured in cecal content from germ-free (GF), easily accessible microbiota (EAM), low-complexity microbiota (LCM), Oligo-MM consortium (OLI), and specific pathogen-free (SPF) mice.

### 2.2 Applications in microbiome research: Quantification of the fecal N-metabolome in human feces and mouse gastrointestinal tract content

Fig. 3A presents a simplified model of nitrogen exchange between the liver and gut bacteria. To test if our approach could measure relevant metabolites in human gut content, we applied our technique to fecal samples from 9 adult human donors. A principal component analysis (PCA) of the amine-containing metabolite concentrations in all samples revealed individuals 52 and 37 as having the most distinct fecal N-metabolomes (Fig. 3B). FST_52 had particularly high ammonia and cadaverine abundances, approximately 61 nmol/mg feces for each compound after normalization (Fig. 3C–D). Ammonia was detected in all nine donors, whereas cadaverine abundance was markedly higher in FST_52 than in the remaining samples. Although this sample size is much too small to link these changes to any host characteristics (diet, medication, pre-existing conditions), we speculate that analyzing fecal nitrogen metabolome is likely to reveal medically relevant changes in host physiology and gut microbiome function. For example, an altered activity of proteolytic bacteria could be a potential cause for altered levels of ammonia and free amino acids in feces without corresponding changes in urea concentration.^49,50^

**Figure 3.**
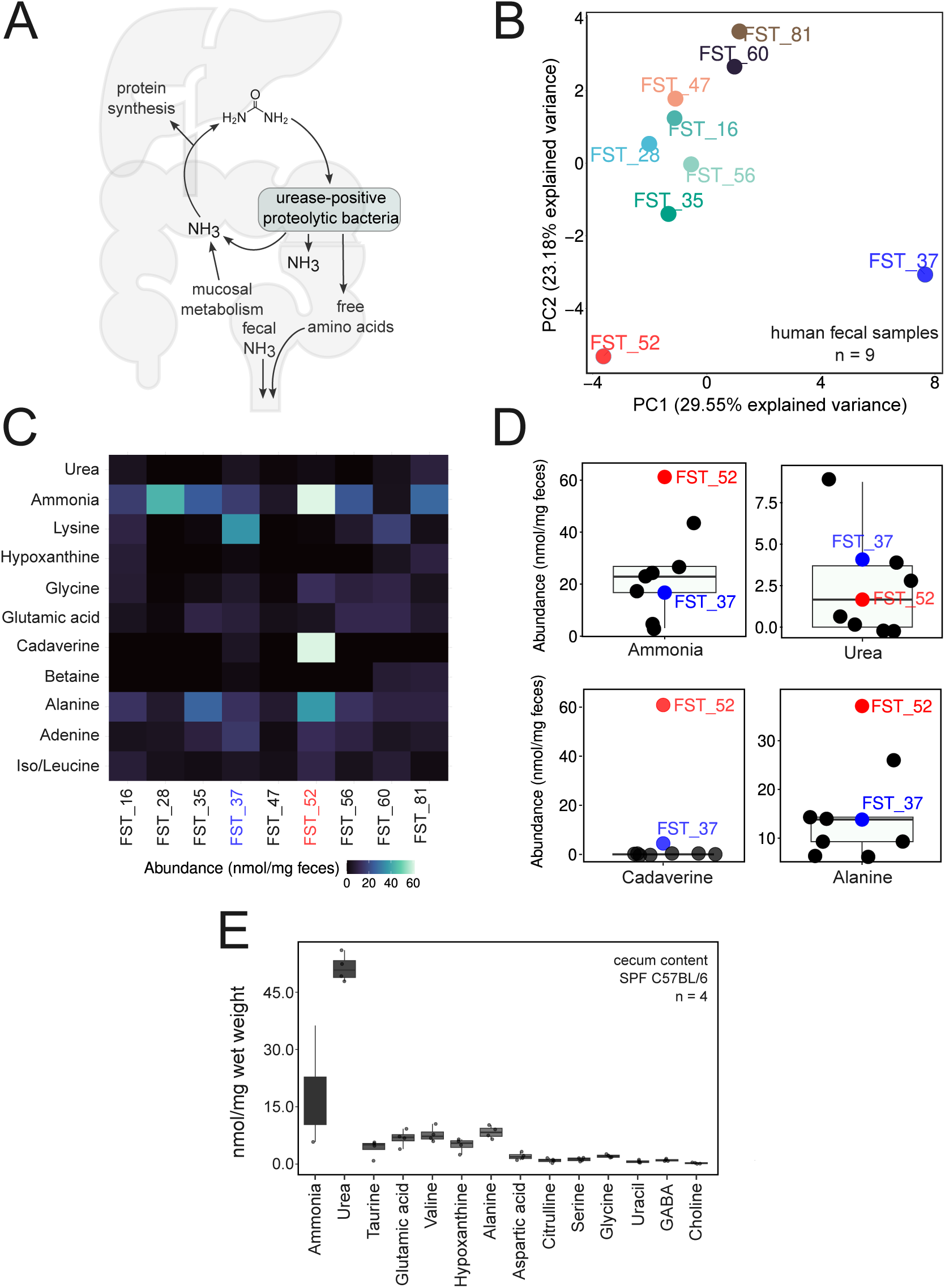
Targeted nitrogen-metabolite profiles in human feces and SPF mouse cecal content. **A)** Model of nitrogen exchange between the gut and liver. **B)** PCA of centered and scaled targeted metabolite abundances from nine adult human donors; each point represents one donor. **C)** Heatmap of 11 selected compounds across the nine donors, expressed as nmol/mg feces. **D)** Boxplots of ammonia, urea, cadaverine, and alanine abundances in the same donors, in nmol/mg feces; FST_52 is highlighted in red and FST_37 in blue. Boxes show the median and interquartile range, whiskers extend to the most extreme observations within 1.5 times the interquartile range, and points show individual donors. **E)** Abundance estimates for 14 selected compounds in cecal content from four SPF C57BL/6 mice, expressed as nmol/mg wet content. Boxes and whiskers are defined as in D; points represent individual mice.

To explore applicability to a common model animal, we examined the murine cecal nitrogen metabolome in laboratory C57BL/6 mice with a specific-pathogen-free microbiome fed standard chow diet. Abundance estimates for selected compounds illustrate the variation among individual mice (Fig. 3E). The presence of variable amounts of some amino acids and ammonia in mouse and human feces is likely due to both host and microbiome metabolism. In comparison to commercial colorimetric assays, where we observed suppression of weak signal by some gut content matrices (Fig. 2), the values obtained using our semi-targeted LC-HRMS method were reproducible between samples, and ammonia concentration are aligned with published values.^48,51–53^

### 2.3 Applications in fundamental microbiology: Quantifying N-assimilation in E. coli as a function of nitrogen source

While many studies are available on the nitrogen regulation systems of *E. coli,* most studies focusing on precise metabolite quantitation (e.g., via LC-HRMS) predominantly target its carbon metabolism pathways.^91–95^ To demonstrate the relevance of our approach for understanding *in vitro* microbial metabolism, we grew the reference *E. coli* strain MG1655 in M9 minimal medium (see Materials and methods) with either glutamine or ammonium chloride as the nitrogen source. This was compared to *E. coli* grown in lysogeny broth (LB), a nutrient-rich medium. In LB, the growth of MG1655 is limited by carbon but not nitrogen availability, and the media contains a range of N-sources including high concentrations of amino acids. Both the bacterial pellet and the spent medium were analyzed to compare biomass-associated and extracellular nitrogen-metabolite profiles (Fig. 4B–E).

**Figure 4.**
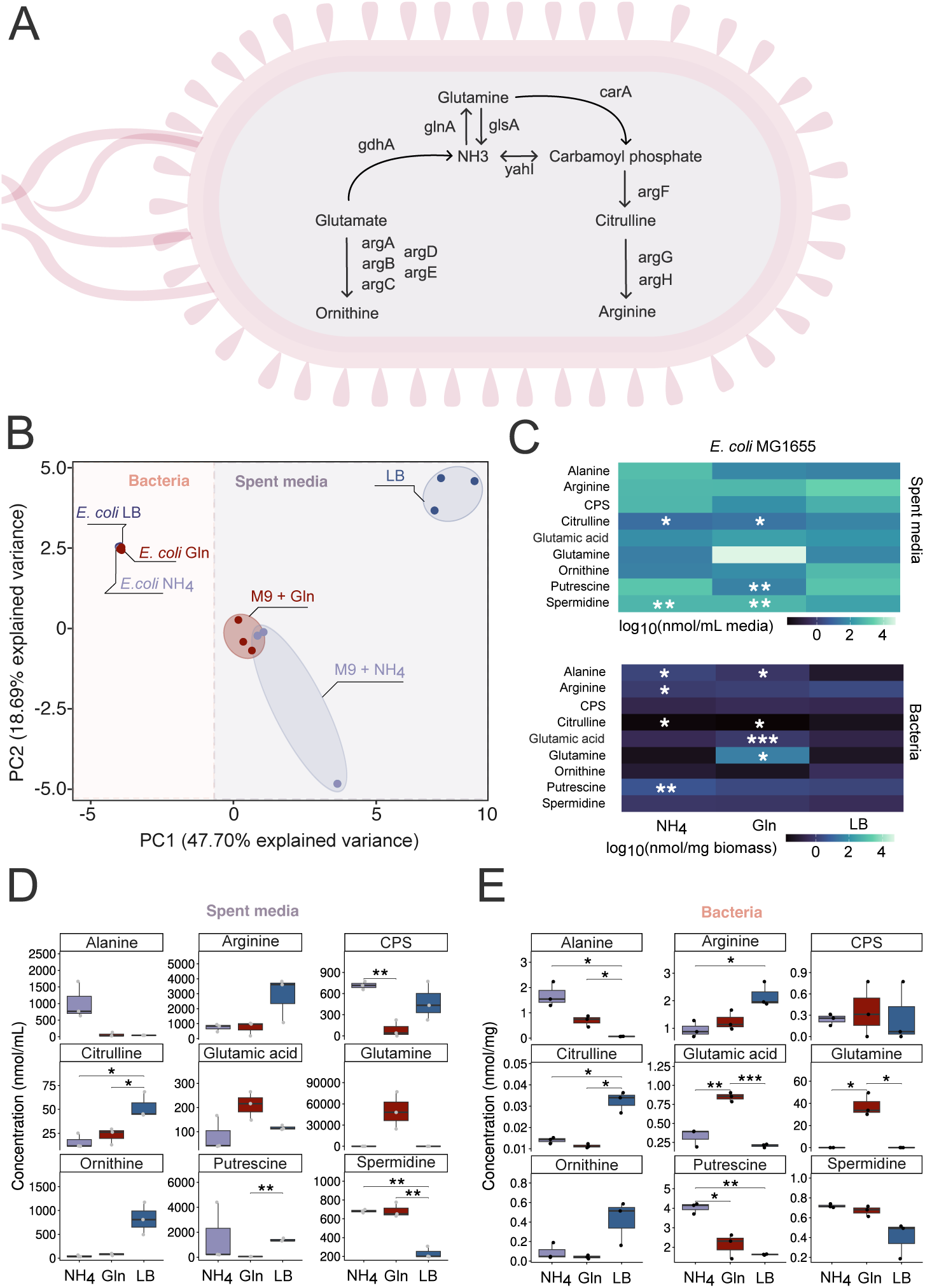
Nitrogen-metabolite profiles of *E. coli* MG1655 grown with different nitrogen sources. **A)** Schematic of nitrogen assimilation and associated glutamate, glutamine, ornithine, and arginine pathways. **B)** PCA of centered and scaled targeted profiles from bacterial pellets and spent media after growth in M9 with ammonium chloride (NH4), M9 with glutamine (Gln), or lysogeny broth (LB). **C)** Heatmaps of nine selected compounds, showing log_10_-transformed group means separately for spent medium (nmol/mL) and bacterial biomass (nmol/mg). Asterisks mark comparisons with LB within each fraction. **D–E)** Boxplots of the same compounds in spent medium (D) and bacterial pellets (E), with three samples per medium and fraction. Points show individual samples; boxes show the median and interquartile range, with whiskers extending to the most extreme observations within 1.5 times the interquartile range. Brackets show pairwise comparisons between media within each fraction. Comparisons in C–E use two-sided Welch’s t-tests with p values: * p ≤ 0.05, ** p ≤ 0.01, *** p ≤ 0.001. CPS = carbamoyl phosphate.

The metabolite profiles differed between bacterial pellets and spent media and among growth conditions (Fig. 4B). Within the pellet fraction, glutamine-fed bacteria had higher glutamine and glutamate abundances than bacteria grown with ammonium chloride, whereas alanine and putrescine were higher in the ammonium-fed group (Fig. 4C,E). LB-grown cultures showed higher citrulline abundance in both fractions, while glutamine accumulated in the spent medium of glutamine-fed cultures (Fig. 4C–E). These measurements demonstrate the application of the method to bacterial biomass and spent medium, where metabolite abundances describe the measured pools and serve as a proxy for nitrogen assimilation.

### 2.4 Applications in environmental ecology: Cnidaria–algae symbiosis

Symbioses between animals and algae/bacteria generate a vast range of ecologically important functions. A prominent example occurs in cnidaria – a group of animals that includes corals and sea anemones – where associations with symbiotic algae are central to the health of coral reefs. As corals are challenging to maintain under lab conditions, this symbiosis is widely studied in the laboratory setting using sea anemones as a model system,^54^ which can be maintained with or without symbiotic algae. Withholding nitrogen is the hypothesized mechanism by which the animal host limits overgrowth of the photosynthetic symbiont, thereby stabilizing the symbiosis.^55–57^ Thus, accurately quantifying nitrogen metabolites is of relevance to advance our understanding of symbiosis maintenance, as well as host-microbe interactions underlying coral bleaching.^58^ In our study, we quantified the nitrogen metabolome of laboratory-cultured sea anemones, with (symbiotic) or without (aposymbiotic) intracellular algae, the surrounding water, as well as algae alone.

Mean ammonia abundance was 2.05 ± 0.50 nmol/mg wet weight in aposymbiotic anemones and 1.16 ± 0.12 nmol/mg wet weight in symbiotic anemones (mean ± SD, n = 3 per group; Fig. 5C), consistent with the hypothesis of nitrogen restriction by symbiotic hosts. Mean abundances of TMAO, proline, (iso)leucine, and valine were also lower in symbiotic anemones, whereas mean citrulline abundance was higher (Fig. 5C). Data-dependent acquisition also allowed us to examine underivatized ions alongside DEEMM derivatives. For example, a sodium adduct assigned to betaine, an important nitrogen reserve produced by coral reef cnidaria,^59^ was detected in anemones and showed higher signal in symbiotic anemones than in the combined free-living algal cultures (Fig. 5D).

**Figure 5.**
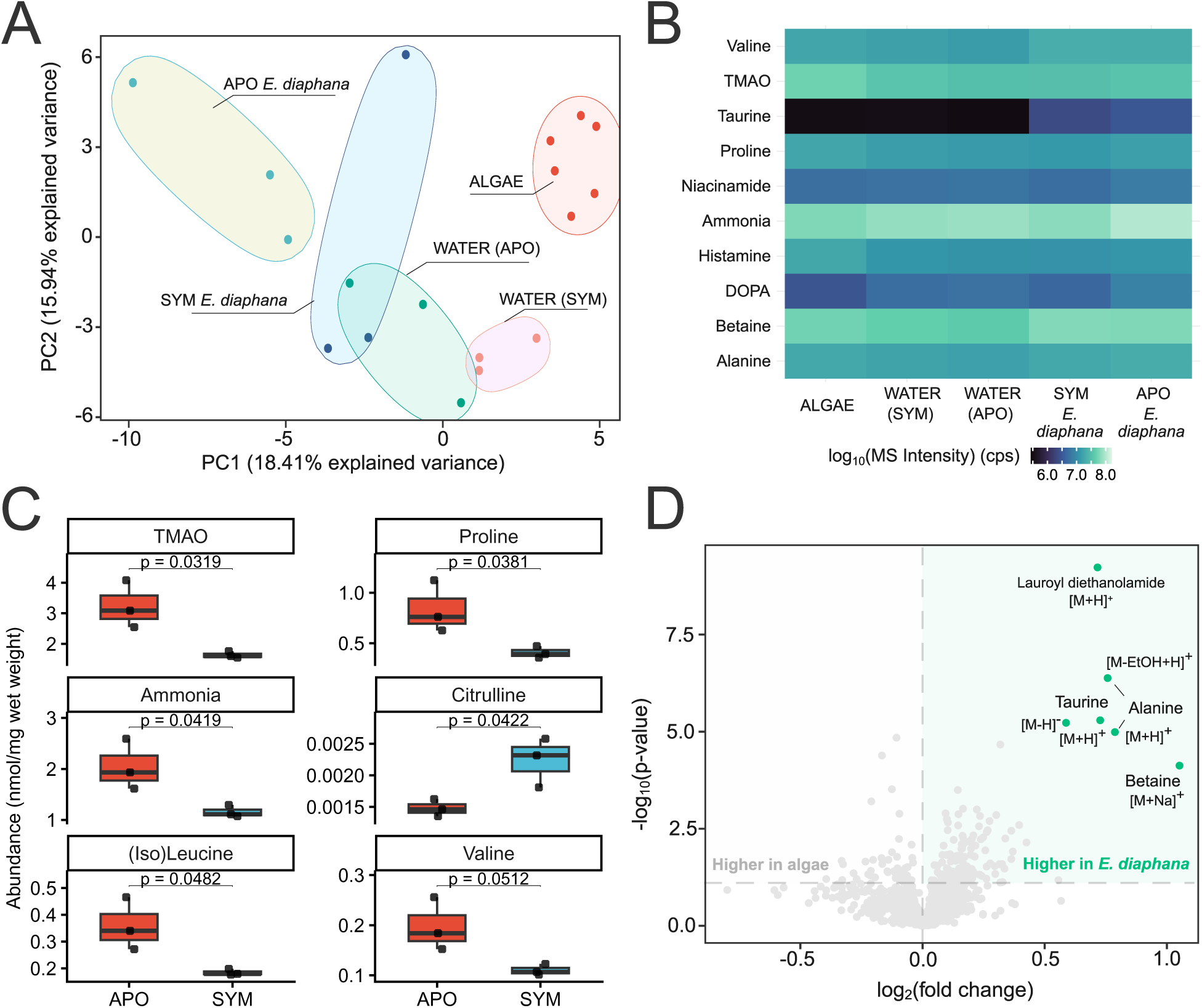
Differences in the nitrogen metabolome of symbiotic and aposymbiotic sea anemones. **A)** Principal component analysis of all m/z features found in samples of symbiotic (SYM) and aposymbiotic (APO) *Exaiptasia diaphana* F003 sea anemones, the surrounding water (WATER), and axenized cultures (bacteria-free) of the algal symbionts *Symbiodinium linuchae* SSA01 and *Breviolum minutum* SSB01 (summed as ALGAE), with three replicates per condition. **B)** Heatmap showing the logarithmic intensities of the ten most variable N-metabolites among the sea anemones, the surrounding water, and the algae. **C)** Boxplots of TMAO, proline, ammonia, citrulline, (iso)leucine, and valine abundances in aposymbiotic (APO, red) and symbiotic (SYM, blue) sea anemones, expressed as nmol/mg wet weight. Points represent individual animals (n = 3 per group). Boxes show the median and interquartile range; whiskers extend to the most extreme observations within 1.5 times the interquartile range. **D)** Volcano plot showing log2 fold change differences between summed algal samples and symbiotic sea anemones, with values on the right indicating higher production in the latter. Statistically significant compounds upregulated in the SYM group are highlighted in green. DOPA = 3,4-dihydroxy-L-phenylalanine; TMAO = trimethylamine oxide. Note that a surfactant contaminant from the preparation generates a significant, but biologically uninteresting hit (marked on the plot).

### 2.5 Applications in environmental ecology: aquatic system diversity and pollution

Nitrogen-containing organic compounds are of interest in wastewater treatment and freshwater analysis.^60–62^ To test the application of DEEMM derivatization and LC-HRMS to aquatic samples, we collected water from three sources in and around Zurich, Switzerland: the west edge of the Katzensee lake, a pond next to it called Büsisee, and the river Limmat about 4 km downstream of the Letten hydropower plant. These were compared with pre-treatment water samples from treatment plants in Bern, Zurich, and Basel. The displayed compounds include ammonia, a routinely monitored water-quality parameter,^19^ and amino acids such as alanine and glycine, which contribute to dissolved organic matter.^63^ Caffeine was included as a marker of anthropogenic input. Targeted panels report calibration-based concentration estimates in nmol/mL of original water. Considering the high dilution of water samples, accurate quantification would require calibration standards covering lower concentrations than our regular range (1 µM to 10 mM). For the untargeted comparison, we compared only raw MS intensities.

On average, we found higher concentrations of ammonia and (iso)leucine in the wastewater samples, whereas glycine, alanine, putrescine, TMAO, and caffeine were higher in the freshwater. Although the fresh and wastewater sample types were evidently different from each other and a control ultrapure Milli-Q water measurement, they did not cluster separately according to the sampling site (Fig. 6A). While the technique requires optimization of standard curves for aquatic samples, the observed differences indicate that the application can be extended.

**Figure 6.**
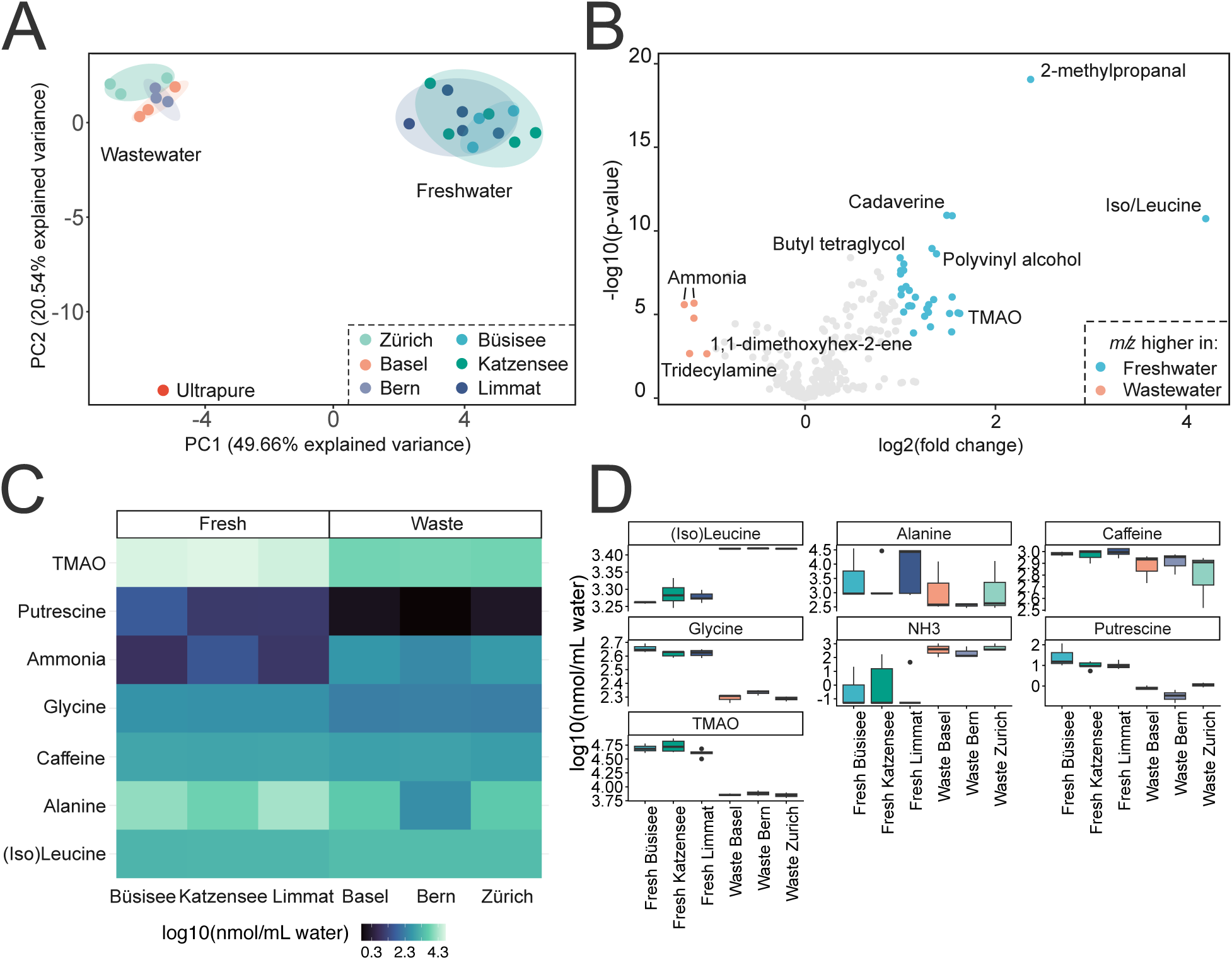
Targeted concentration estimates and untargeted MS profiles in freshwater and wastewater. **A)** PCA of targeted profiles from Büsisee (n = 3), Katzensee (n = 5), Limmat (n = 5), and wastewater from Basel, Bern, and Zurich (n = 3 per city), with an ultrapure-water control. Points represent assayed samples and are colored by source. **B)** Exploratory volcano plot comparing freshwater from Limmat (n = 5) with wastewater from Basel, Bern, and Zurich (n = 9 in total). Two-sided Welch’s t-tests were applied to log_2_ MS intensities. Features with p < 0.05 and absolute log_2_ fold change > 1 are highlighted: positive values indicate higher signal in Limmat (blue), and negative values indicate higher signal in wastewater (orange). Labels indicate putative feature assignments. **C)** Heatmap of group-mean concentration estimates for seven selected compounds: ammonia, TMAO, alanine, glycine, putrescine, caffeine, and (iso)leucine. **D)** Boxplots of the same compounds by sampling source. C–D use log_10_-transformed estimates in nmol/mL of original water. Boxes show the median and interquartile range; whiskers extend to the most extreme observations within 1.5 times the interquartile range, and points beyond the whiskers are outliers. TMAO = trimethylamine oxide.

### 2.6 Applying amine analysis to a biological question: Ammonia and bioactive amines reveal the major role of microbiota in modulating intestinal nitrogen handling

Finally, we applied our method to address a major biologically relevant question: how does microbiome composition influence the concentrations of ammonia and bioactive amines in the gut content? We quantified the bioactive amine and ammonia concentrations in cecum content from “specific-pathogen-free” (SPF) mice, three populations of gnotobiotic mice: low-complexity microbiota (LCM) generated via weak diversification of altered Schaedler flora,^64^ OligoMM12 mice, colonized from birth with a known composition of 12 bacterial species,^65–67^ and “easily-accessible microbiota (EAM)” – colonized mice, consisting of *Bacteroides thetaiotaomicron, Eubacterium rectale* and *Escherichia coli*;^68^ and mice lacking any detectable microbiota (germ-free, GF) (Fig. 7A). The dataset comprised five mice each in the GF, EAM, LCM, and OligoMM12 groups and four SPF mice. All mice had been bred with their respective gut microbial compositions^69–71^ and were fed identical sterile food and water prior to sampling. All animals were sampled in the morning of a standard night-day cycle to avoid circadian rhythm-driven shifts between groups.

**Figure 7.**
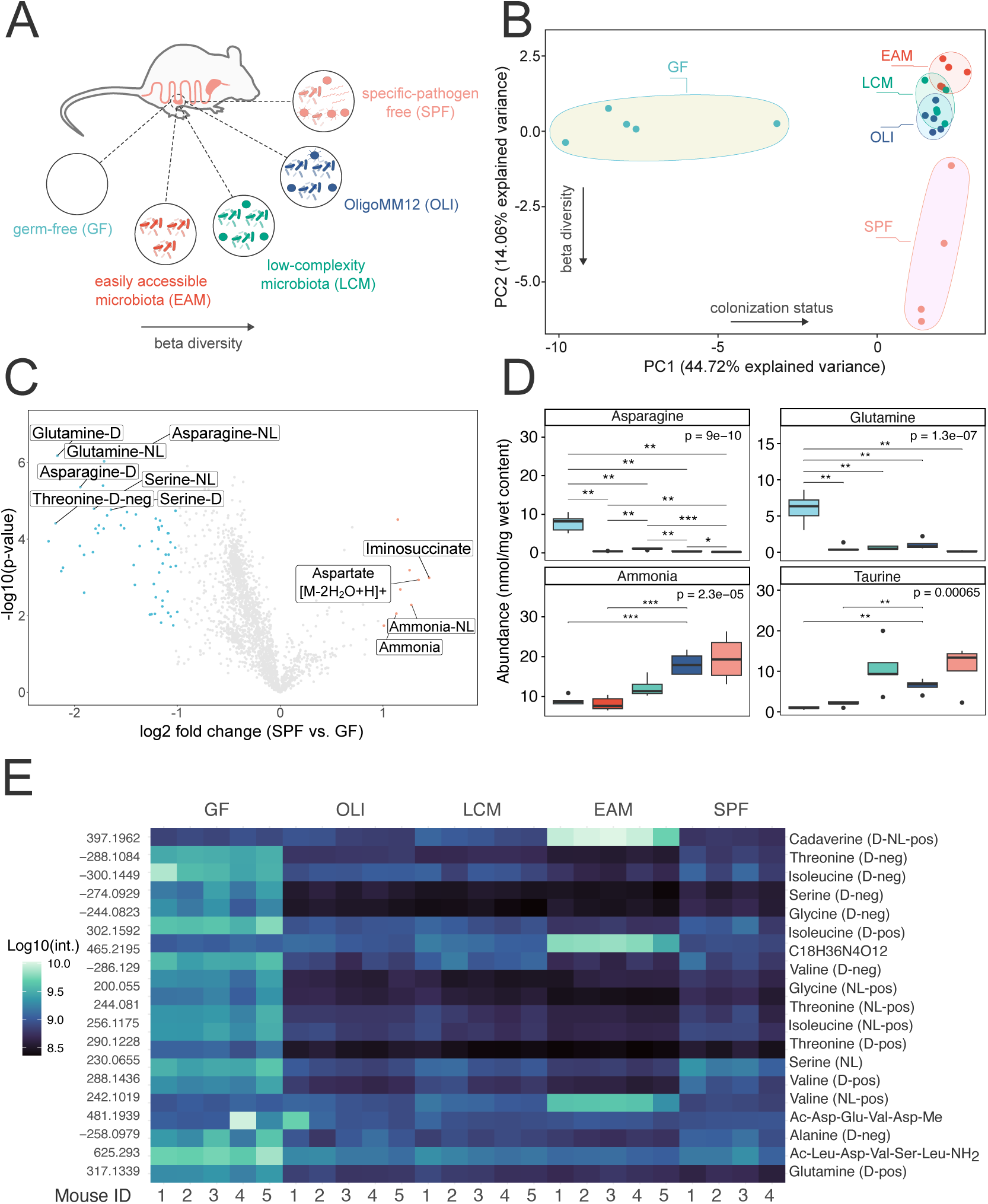
Cecal nitrogen-metabolite profiles in mice with different gut microbiota. The study includes five mice each with germ-free (GF), easily accessible microbiota (EAM), low-complexity microbiota (LCM), or OligoMM12 (OLI) status and four specific-pathogen-free (SPF) mice. **A)** Schematic of the colonization groups. **B)** PCA of centered and scaled untargeted MS features; each point represents one mouse. **C)** Volcano plot comparing SPF with GF mice using two-sided Welch’s t-tests on log_2_ MS intensities. Negative fold changes indicate higher signals in GF, and positive fold changes indicate higher signals in SPF. Features with nominal p < 0.05 and absolute log_2_ fold change > 1 are highlighted. **D)** Boxplots of asparagine, glutamine, ammonia, and taurine abundance estimates in nmol/mg wet cecal content. Boxes show the median and interquartile range; whiskers extend to the most extreme observations within 1.5 times the interquartile range, and dots show outliers. Numeric p values are from one-way ANOVA across the five groups; brackets display pairwise Welch’s t-test p values: * p ≤ 0.05, ** p ≤ 0.01, *** p ≤ 0.001, **** p ≤ 0.0001. **E)** Heatmap of 19 selected variable MS features, colored by log_10_(intensity + 1), with one column per mouse. Labels give m/z values and putative assignments; negative m/z labels denote negative ion mode measurements. D = derivatized ion; NL = neutral-loss ion; pos/neg = ion polarity.

Opting for high-resolution data-dependent acquisition (DDA) over traditional targeted mass spectrometry methods, such as MRM or PRM, enabled us to detect biological fluctuations beyond our target compound list. This was particularly advantageous for this proof-of-concept study, as gut microbial colonization is known to profoundly impact the metabolic status in mice.^72–74^

In the PCA, PC1 separated germ-free from colonized animals, while PC2 distinguished SPF mice from the defined microbiota groups (Fig. 7B). The untargeted comparison showed higher signals for several amino-acid derivatives in GF mice, including glutamine, asparagine, serine, and threonine (Fig. 7C). In the targeted plots, mean asparagine and glutamine abundance estimates were more than ten-fold higher in GF than SPF mice, whereas ammonia and taurine were lower in GF mice (Fig. 7D). It should be noted, that the cecum of a germ-free mouse also has a roughly ten-fold higher volume than that of an SPF-colonized animal due to water retention, so the overall abundance of these amino acids in the large intestine of a germ-free animal is around 100-fold higher than in a colonized mouse. The biological implications of these very high amino acid exposures have not been extensively investigated.

Beyond the target aminoenones, features putatively assigned to iminosuccinate and an aspartate fragmentation product with double water loss showed higher signals in SPF mice (Fig. 7C). The heatmap also showed distinct profiles among the defined microbiota groups that differed from both GF and SPF mice, including a higher cadaverine-associated signal in EAM mice (Fig. 7E), potentially reflecting altered amine metabolism associated with their reduced microbial complexity. Most, but not all, nitrogen metabolite concentrations differed between GF and colonized mice. Features assigned to short peptides by MS2 showed group-dependent intensities, with some enriched in GF or EAM mice but not in mice with more diverse microbiota, potentially reflecting altered digestion or production of these peptides. While some of these differences have previously been reported by untargeted metabolomics, quantification reveals their extent and warrants exploration of the health consequences of high amine exposure. Together, the targeted abundance estimates and untargeted MS profiles show a substantial association between gut microbiota composition and the cecal nitrogen metabolome, motivating further investigation of the physiological consequences.

## 3 Discussion

Tracing nitrogen containing compounds and particularly ammonia in complex biological matrices, while of great implication in several fields of biology and environmental sciences, has been technically challenging.^75–79^ There was a lack of reliable and reproducible methods which provide absolute quantification of a large set of nitrogenous metabolites and are applicable to a wide range of possible samples.^80–82^ Therefore, we combined existing approaches with a dedicated analytical pipeline to design a semi-targeted mass spectrometric method for accurate detection and quantification of over 50 compounds pertaining to nitrogen metabolism, and, as proof of concept, applied it in different biologically relevant environments.

A recent study by Maciel et al.^35^ proposed the usage of neutral loss mode (NLS) for mass spectrometric analysis of DEEMM-derivatives, as they exhibit a characteristic 46 Da fragmentation pattern upon being subjected to gas-phase collision or high voltage (Fig. 1B). While this approach is highly effective for isolating all DEEMM aminoenones, it does not include the other potentially interesting metabolites present in the sample matrix. Additionally, relying exclusively on DEEMM derivatives may lead to less precise quantification, as conversion is not always complete after the derivatization reaction, and some primary amines do not react at all. Because of its selectivity, DDA with high-resolution mass spectrometry is advantageous in discovery workflows and allows for confident quantification of the abundance of ions contributing to the total concentration of the analyte.

The main nitrogen-containing small molecules in fecal matter are amines, amino acids, nucleosides, amino ketones, and indole derivatives. Polyamines and ammonia contribute important information about the gut, such as microbial activity, metabolism, and cell turnover.^83–85^ In human stool samples, we observed one healthy volunteer who was an outlier in terms of ammonia content (Fig. 3B-D). While we cannot speculate on mechanisms based on the very small sample size and limited metadata, the very high concentrations of ammonia and cadaverine in this individual could be health relevant, and understanding this variation at a population level could reveal important microbiome-diet-health links. Polyamines such as spermidine and cadaverine have been linked to colon cancer, inflammatory bowel disease, ulcerative colitis, but also anti-inflammatory properties and maintenance of mucosal homeostasis. The LC-HRMS approach presented here, with accompanying software solutions,^45^ is broadly accessible and will allow clinical studies to test these links.

Nitrogen compounds play an important role in many symbioses, often involving bacteria that fix molecular nitrogen, recycle organic nitrogen waste, or re-oxidize inorganic ammonium for their animal, plant or algal hosts. In the cnidaria-algae symbiosis, animal hosts withdraw intracellular nitrogen from their algal symbionts by locking it up in N-compounds, such as non-essential amino acids^86^ or dipeptides,^87^ which stimulates the release of photosynthetically fixed carbon from the algae to the host.^58^ Our method can reveal details of this symbiosis by quantifying the concentration of N-containing compounds in a single low-volume sample, while being applicable to samples with high salt concentration. In line with biological expectations, we confirmed lower ammonium concentrations in symbiotic anemones, likely reflecting the assimilation of ammonium via the glutamine synthase (GS/GOGAT pathway) by the host. Further, reduced concentrations of taurine were detected, which is a “host release factor” that can tune photosynthate excretion in algae.^88,89^ We could also detect phenolic DOPA and methylated amine betaine relevant for wounding responses and nitrogen storage, indicating potential to expand the assay for this particular application. We are confident that our method of N-metabolite profiling can be developed to monitor the state of the cnidaria-algae symbiosis, for example in coral reefs.

We also show preliminary evidence that the technique can be applied to fresh water monitoring. While fresh water is a simple matrix and colorimetric assays for ammonia are typically reliable, the ability to understand biological nitrogen flux in acquatic systems can allow more detailed monitoring and interventions. For example, elevated urea concentrations are a result of increased breakdown of organic matter and excretion by marine organisms, or a product of agricultural runoff and detailed quantification may differentiate these causes. Although further work is required to optimize standard concentrations for appropriate calibration, our semi-targeted LC-HRMS method could be applicable in this field.

Finally, we applied amine analysis to understand the influence of the gut microbiome on amine abundance in the murine large intestine. Gnotobiotic animals, i.e., germ-free animals that have been selectively colonized with known microbial populations, have generated valuable insights into how the gut microbiome influences host health over the past 20 years.^90^ While a lot of work has been done on carbon metabolism of microbiomes,^69,91–95^ we still know relatively little about nitrogen metabolism of these consortia. Elevated amino acids and decreased ammonia levels in the gut of germ-free mice versus colonized animals has been observed in semi-quantitative analysis.^48,96,97^ Here we could accurately quantify these differences and show that in many cases the concentration of amino acids is more than ten-fold elevated under germ-free conditions. It should be noted that as the volume of gut content can also be ten-fold higher in germ-free mice, the overall abundance of these compounds in the germ-free large intestine is potentially 100-fold higher than in and SPF-colonized mouse. Interestingly, we observe that gnotobiotic colonizations typically generate intermediate phenotypes. Even very simple microbiota seem to be sufficient to decrease the concentration of most amino acids to the levels observed in colonized animals. However, mice colonized only with *E. coli, Bacteroides thetaiotaomicron and Agathobacter rectalis* presented very high levels of cadaverine in cecal content, which is not observed in either germ-free or other colonized animals and must be a produce of the very limited metabolic capacity of this minimal microbiome. As the LC-HRMS technique also works reliably on bacteria and supernatant of *in vitro* cultures in minimal and complex culture media, there is considerable potential to determine individual species contributions to the overall N-metabolome and interesting follow-up studies could explore the influence of different durations, classes and dosing of antibiotics on gut nitrogen metabolomes. The ability to accurately quantify the N-metabolome in gut content and tissues of experimental animals opens this field to detailed quantitative analysis, with potential to understand the influence of gut microbiome nitrogen metabolism on host health. We propose that this is an under-explored area of host-microbiome interactions which has been limited by poor resolution of the available measurement techniques.

## 4 Conclusion

In summary, we described an analytical mass spectrometric method based on DEEMM-derivatization for the detection and quantification of 50 compounds comprising the nitrogen metabolome, and present multiple applications of it in ecologically diverse biological matrices. Owing to good chromatographic resolution, we are able to precisely quantify compounds without overbearing any of the components of the mass spectrometer with an influx of competing analyte ions. The method is compatible with multiple-well plate-based workflows and requires runtimes of 30 minutes, making it viable for moderate to high-throughput analyses. The limitations include relatively slow sample preparation and the need for derivatization, which is not compatible with all analytes of interest. The latter can be addressed by implementing our data acquisition and analysis pipeline, utilizing DDA and the open-source tool for targeted MS analysis, LCMSpector.^45^ Some applications of the method will need modifications of the protocol, for example to remove excessive contaminants and interferants, or change the concentration range of standard curves to better suit the expected sample content. Nonetheless, we believe this method is broadly applicable to very many areas relying on precise and repeatable quantification of nitrogen-containing compounds, is broadly implementable wherever LC-HRMS equipment is accessible and has potential to radically improve the quantification of these molecules.

## 5 Materials and methods

### 5.1 Mouse experiments

All animal experiments were conducted according to legal and ethical regulations and approved by the Swiss Cantonal authorities (License ZH010/24, ZH184/24).

Germ-free, gnotobiotic (EAM, Oligo-MM12, LCM) and Specific Pathogen Free (SPF, complex microbiota) C57BL/6 wild type mice were bred and maintained at the ETH Phenomics Center on a standard diet in individually ventilated cages under strict hygiene conditions. For organ collection, animals were euthanized by CO_2_ asphyxiation and exsanguination at 8-14 weeks of age. After removing the whole gastrointestinal tract for tissue collection, cecum content was collected, immediately snap frozen in liquid nitrogen, and subsequently stored at −80°C until further processing.

### 5.2 Chemical reagents

Standards of all the traced amines and amino acids were purchased from Sigma Aldrich unless otherwise specified.

#### Extraction solutions

Perchloric acid analytical grade (70%), Sigma-Aldrich, cat. no. 30755-500ml Hydrochloric acid, analytical grade, VWR International, cat. no. VWRC20252.290-1lt.

#### Derivatization reaction reagents

DEEMM (diethyl-ethoxymethylenemalonate), VWR International, cat. no. TCE0255. Boric acid, Sigma-Aldrich, cat. no. 31146-500G.

#### LC-MS solutions

Formic acid (FA), LC/MS Ultra, >98% purity; Sigma-Aldrich, cat. no. 14265-1ML.

Water, Optima LC/MS Grade, Fisher Chemical, cat. no. 10505904-2.5L.

Methanol, Optima LC/MS Grade, Fisher Chemical, cat. no. 10767665-2.5L.

Ammonium acetate, >98% purity; Sigma-Aldrich, cat. no. A7262-500G.

Acetonitrile, Optima LC/MS Grade, Fisher Chemical, cat. no.10001334-2.5L.

### 5.3 Buffer solutions

Maintaining the precision and accuracy of chromatographic separation steps and limiting contamination of high-resolution mass analyzers requires all solutions to be prepared using LC-MS grade solvents and reagents. Before usage, buffer solutions are degassed and filtered using 0.45 μm PTFE membrane filters. To ensure the best chromatographic performance, buffers should be prepared fresh before analysis and filtered daily to avoid microbial growth.

### 5.4 Equipment

High mass accuracy and high resolving power are necessary for reliable DDA experiments and can substantially influence subsequent data analysis steps (e.g., LCMSpector, SIRIUS). Fourier-transform mass analyzers (Orbitrap or FT-ICR) are most suitable for this purpose.

Additionally, for high-throughput analysis, the scan speed of the mass analyzer needs to be considered. To obtain sufficient sampling points per chromatographic peak at high MS and MS/MS resolution (>100’000), the DDA scanning needs to be as quick as possible, which necessitates the use of state-of-the-art spectrometers.

### 5.5 Batch design

While the method is fully compatible with high-throughput workflows, the high salt concentrations and potential sample matrix influence (especially when analyzing gastrointestinal tract content) will rapidly contaminate the ion source, as well as the sensitive ion optic parts of the mass spectrometer and consequently degrade its performance over time. Therefore, the batch design necessitates the use of frequent cleaning steps. We recommend running a cleaning step followed by a blank control injection every five measurements. If the number of samples in a batch is high (e.g., >100), it is recommended to perform more thorough cleaning every ten samples.

After particularly contaminating sample batches, such as those containing highly viscous, oily or otherwise water-insoluble analyte solutions or non-removable particulates, one to several isocratic wash runs of 0.1% FA (LC/MS ultra grade) from the LC system is performed at a flow rate of 200 μl × min^−1^ for 60 minutes, and the changes in system pressure is monitored closely over the course of washing.

### 5.6 Data acquisition by LC-HRMS

Samples were injected into a Vanquish Flex (ThermoFisher Scientific, Germany) liquid chromatograph coupled to a heated electrospray ion source (H-ESI) of an Orbitrap Exploris 240 mass spectrometer (ThermoFisher Scientific, Germany). Reverse-phase chromatography was run on a 1.7 μm (2.1 mm × 100 mm) Acquity BEH C18 UPLC column (Waters, Switzerland) operated at 40°C, with a dedicated Vanguard C18 column guard attached. The mobile phase was a gradient mixture of aqueous 10 mM ammonium acetate, water, methanol, and acetonitrile. Detailed information on the gradient can be found in the Supporting Information. Absorbance data for each run were collected at 230 nm and 280 nm using the built-in Vanquish Flex diode array detector (Supporting information – Fig. S2C). The Orbitrap mass spectrometer was operated in both the positive and negative ionization modes, each recorded in a separate run for all analyzed samples. In both modes, the sheath gas was set to 30 a.u., the auxiliary nebulizer gas to 10 a.u at 400 °C, and the ion transfer tube was heated to 350 °C. In positive mode, the capillary voltage was set to 3.4 kV, and in negative mode to 2 kV. Data were acquired as full-scan MS1 recorded in profile mode, followed by a series of 10 consecutive data-dependent MS2 scans recorded in centroid mode, with dynamic exclusion filtering applied to every 60-second acquisition window.

### 5.7 Data processing

Raw MS data files in the proprietary ThermoFisher Scientific RAW file format were first converted into the open mzML format using MSConvert (ProteoWizard package).^98^ Raw LC chromatograms were directly exported from the Chromeleon Chromatography Data System in the .txt file format. The mzML and .txt files were subsequently analyzed using LC-Inspector, a standalone user-interface application for viewing and analyzing LC-MS data.^99^ The processed .csv files, containing all the targeted primary ions, their absorbance values and MS intensities, as well as concentrations extrapolated from calibration curves of standard samples, were then exported and further analyzed with R. The study data and analysis code are available in Zenodo (10.5281/zenodo.22881185).

Data-dependent acquisition tandem mass spectra (DDA-MS2) were processed and annotated with SIRIUS.^100^ Detailed information containing the putative assignments of fragment ions can be found in the Supporting Information.

### 5.8 Sample processing

Metabolites were extracted with a clean metal bead by high-speed disruption (TissueLyzer, Qiagen) at 25 Hz for 3 minutes in a three-fold excess (v/w) of 100 mM perchloric acid, using pre-cooled 2 mL Eppendorf tube adaptors to minimize the effect of heat generation during shaking. The homogenized solution was centrifuged (20 minutes, 4°C, 6 000 g), and upon transferring to a new tube, centrifuged again (20 minutes, 4°C, 14 000 g). The resulting supernatant was protein precipitated by adding 100 μl of cold methanol, incubating on ice for 10 minutes, shaking (3 minutes, 4°C, 2 000 rpm), and centrifuging (15 minutes, 4°C, 5 500 g). The supernatant was then frozen or processed further by derivatization.

### 5.9 Amine derivatization with diethyl ethoxymethylenemalonate (DEEMM)

The derivatization reaction was carried out on previously homogenized and protein-precipitated samples.^41^ 37.5 μl of the processed sample was mixed with 141.25 μl of a derivatization mix (87.5 μl of 1 M orthoboric acid, 37.5 μl of 0.1 M hydrochloric acid, 12.5 μl of methanol, 2 μl of 2 g/l aminoadipic acid, 1.75 μl of diethyl ethoxymethylenemalonate) and incubated for 2 hours at 70°C. The samples were then syringe-filtered (0.22 μm) into HPLC vial inserts and analyzed.

### 5.10 Colorimetric assays

Cecum content samples were initially diluted 1:15 w/v with ddH_2_O and homogenized for 3 minutes at 25 Hz in a QUIAGEN TissueLyzer II with a stainless-steel bead per tube. Samples were then briefly spun at 100 x g for 30 seconds to deposit food debris and supernatant was used for measurements according to manufacturer’s instructions (B: Biorbyt, orb545637; S: Sigma, MAK310).

### 5.11 Bacterial culture and media preparation

*Escherichia coli* MG1655 was streaked from glycerol stock onto LB agar plates and incubated overnight at 37°C. Single colonies were picked and used to inoculate pre-cultures in one of the following media: (i) lysogeny Broth (LB); (ii) M9 minimal medium supplemented with 10 mM NH_4_Cl; (iii) M9 minimal medium supplemented with 5 mM L-glutamine. Pre-cultures were grown aerobically at 37°C with shaking. Growth was monitored using a Tecan plate reader, and growth curves indicated that cultures remained in exponential phase until approximately 16–18 h (data not shown). For each condition, biological replicate cultures were established by inoculating fresh medium with 1 µL of the respective pre-culture and incubating at 37°C with shaking for another 16 h.

M9 mineral medium was prepared using a 10× M9 salt solution and sterile water. The final 1× M9 medium contained: Na_2_HPO_4_ (33.7 mM); KH_2_PO_4_ (22.0 mM); NaCl (8.55 mM); Glucose (0.4% w/v); MgSO_4_ (1 mM); CaCl_2_ (0.3 mM); Biotin (1 mg/L); Thiamine (1 mg/L); supplemented with 10 mL of a 100× trace elements solution composed of EDTA (13.4 mM); FeCl_3_·6H_2_O (3.1 mM); ZnCl_2_ (0.62 mM); CuCl_2_·2H_2_O (76 µM); CoCl_2_·2H_2_O (42 µM); H_3_BO_3_ (162 µM); MnCl_2_·4H_2_O (8.1 µM). The pH was adjusted to 7.2 prior to sterilization where applicable. Heat-stable components were autoclaved, while heat-labile components (e.g., vitamins and trace elements) were filter-sterilized (0.22 µm) and added after cooling. For nitrogen-source comparisons, M9 medium was supplemented with either: NH_4_Cl (final concentration 10 mM) or L-glutamine (final concentration 5 mM).

Optical density at 600 nm (OD_600_) was measured and culture volumes corresponding to a total of 5 OD units (approximately 5 × 10^8^ cells) were collected for each sample. The required volume was calculated individually based on measured OD_600_ values. Cultures were centrifuged at 5000 × g for 10 min, after which both the cell pellet and the supernatant were carefully collected and processed separately for downstream analysis. Unless otherwise stated, metabolite extraction, derivatization, and LC-HRMS acquisition were performed as described above.

### 5.12 Sea anemone and algal samples

Symbiotic sea anemones (*Exaiptasia diaphana*, strain F003;^101^ obtained from Annika Guse Lab; LMU Munich) were reared in artificial seawater medium (ASW; Tropic Marin Pro-Reef sea salts; 34 ppt salinity and pH 8.2) at 26 °C with 12:12 h light-dark cycle, 30 µmol m^−2^ s^−1^ light intensity, and weekly feeding with *Artemia* AF brine shrimps (SEP-*Art*, INVE Aquaculture). Aposymbiotic F003 sea anemones were maintained in darkness under the same condition but adjusted to the diurnal light regime 10 weeks before sampling. Three individual specimens per condition (symbiotic: 20 ± 8 mg wet weight; aposymbiotc: 25 ± 8 mg wet weight; maintained in separate rearing boxes) were sampled for N-metabolome analysis six days after feeding, together with 500 μl of the surrounding water, by plunging the samples into liquid nitrogen. The algal strains *Symbiodinium linuchae* SSA01^102^ and *Breviolum minutum* SSB01^103^ both obtained from Annika Guse Lab, LMU Munich), which are endogenous to *Exaiptasia diaphana* F003, were cultured under the same light and temperature regime, but with additional Daigo’s IMK supplements (0.252 g/L; FUJIFILM Wako Chemicals) added to the ASW medium. Algae were axenized by transferring single colonies grown on ASW+IMK agar plates with KAS antibiotics (34 g/L Tropic Marin Pro-Reef sea salts, 12 g/L BD DIFCO™ Noble Agar, 100 μg/mL Ampicillin, 50 μg/mL Streptomycin, 50 μg/mL Kanamycin; adapted from^103^ for five passages on fresh agar plates for one year. For N-metabolome analysis, replicate culture flasks (n = 3 per strain) containing 200 mL ASW+IMK liquid medium (without KAS) were inoculated with algae from agar plates and cultivated for 52 days under the above temperature and light regime. For N-metabolome sampling, 500 μl cultures were taken and rapidly frozen in liquid nitrogen. In parallel, algal cell numbers were determined by flow cytometry (SSA01: 842,167 ± 90,689 cells mL^−1^; SSB01: 337,983 ± 18,820 cells mL^−1^), and the absence of culturable bacteria confirmed by plating on 1/10 MB agar medium (3.74 g/L BD Difco Marine Broth 2216, 30.6 g/L Pro-Reef sea salts, 12 g/L BD Difco Noble Agar; no colonies formed within 4 weeks of incubation at 26 °C in light).

### 5.13 Freshwater samples

To generate the most representative sample possible for the given lake, lentic fresh water samples were collected at different depths in at least 3 different locations, as still-water bodies are unlikely to be homogenous.^104^ The water was sampled using an in-house sampling device consisting of a simple siphon pump attached to a Falcon test tube, into which at least 10 ml of sample was collected. Lotic samples (from the river Limmat) were sampled in multiple spots, and effort was made to sample only from sites with unidirectional flow to avoid mixing streams.

### 5.14 Human fecal samples

The human fecal samples were collected as part of a research study involving persons, title: “Intestinal microbiome strain-level stability in NEC/ or LOS patients” (submitted to the Zurich Cantonal Ethics Comission, BASEC number 2020-01764, ClinicalTrials.gov ID: NCT04792918). Information on gastrointestinal symptoms and antibiotic intake was additionally collected from participating individuals. The participants were instructed to collect the samples into sampling tubes which were then transferred to the processing facility within 2 hours and subsequently flash frozen. Samples for N-compound analysis were taken from frozen feces material and weighed to obtain at least 50 mg of weight and subsequently processed according to the procedure described above.

## Author contributions

MF, ECB, and ES designed the study. MF designed the LC-HRMS method and data analysis pipeline. ECB designed the sample preparation protocol. ECB performed the colorimetric assay ammonia measurements. JK, ECB, and AH carried out the mouse experiments. MS performed the sea anemone cultivation and sampling. ECB and EH carried out sample preparation. MF, ECB, and EH carried out water sampling. MF and EH performed the LC-HRMS experiments. MF and EH analyzed the data. JS participated in the initial LC measurements and data analysis. RZ, SS, and ES reviewed the manuscript. The manuscript was written by contributions of all authors.

## Supporting information

**Table S1.** List of compounds, their m/z values, and names of all the ions used in the method described in this article.

**Figure S1.** Evaluation of colorimetric ammonia quantification assays and sample preparation effects.

**Figure S2.** LC gradient optimization and chromatographic separation of DEEMM-derivatized nitrogen metabolites.

**Table S2.** Elution gradient used for LC separation of DEEMM-derivatized nitrogen metabolites.

**Table S3.** Washing gradient used to clean and re-equilibrate the column between analytical runs.

**Figure S3.** Quantification of nitrogen-containing metabolites in freshwater samples.

## Acknowledgments

We thank Benoit Pugin for their input in sample preparation and designing the gradient used in the initial method based on UPLC. We also thank Prof. Dr. Alexander Harms and the Laboratory of Molecular Phage Biology for providing the non-treated water samples from water treatment plants around Switzerland. S.S. acknowledges core funding from ETH Zürich.

