## Supporting Information for "Comprehensive analysis of ammonia and biogenic amines in ecologically diverse systems with high-resolution mass spectrometry"

### Table of Contents

Table S1. List of compounds, their m/z values, and names of all the ions used in the method described in this article.

Figure S1. Evaluation of colorimetric ammonia quantification assays and sample preparation effects.

Figure S2. LC gradient optimization and chromatographic separation of DEEMM-derivatized nitrogen metabolites.

Table S2. Elution gradient used for LC separation of DEEMM-derivatized nitrogen metabolites.

Table S3. Washing gradient used to clean and re-equilibrate the column between analytical runs.

Figure S3. Quantification of nitrogen-containing metabolites in freshwater samples.

**Table S1.** List of compounds, their m/z values, and names of all the ions used in the method described in this article.

| Compound | m/z values | Ion name |
|---|---|---|
| Adenine | 306.1197 | Adenine-D |
| | 260.0779 | Adenine-NL |
| | 304.1051 | Adenine-D-neg |
| | 136.0618 | Adenine-I-pos |
| | 134.0472 | Adenine-I-neg |
| Adenosine | 438.162 | Adenosine-D |
| | 392.1201 | Adenosine-NL |
| | 436.1473 | Adenosine-D-neg |
| | 268.1041 | Adenosine-I-pos |
| | 266.0894 | Adenosine-I-neg |
| Agmatine | 301.1871 | Agmatine-D |
| | 255.1452 | Agmatine-NL |
| | 471.245 | Agmatine-D-D |
| | 425.2031 | Agmatine-D-NL |
| | 299.1724 | Agmatine-D-neg |
| | 129.1145 | Agmatine-I-pos |
| | 131.1292 | Agmatine-I-neg |
| Alanine | 260.1129 | Alanine-D |

| Compound | m/z values | Ion name |
| --- | --- | --- |
| Alanine | 214.071 | Alanine-NL |
| Alanine | 258.0983 | Alanine-D-neg |
| Alanine | 90.055 | Alanine-I-pos |
| Alanine | 88.0404 | Alanine-I-neg |
| Arginine | 345.1769 | Arginine-D |
| Arginine | 299.135 | Arginine-NL |
| Arginine | 515.2348 | Arginine-D-D |
| Arginine | 469.193 | Arginine-D-NL |
| Arginine | 343.1623 | Arginine-D-neg |
| Arginine | 175.119 | Arginine-I-pos |
| Arginine | 173.1044 | Arginine-I-neg |
| Asparagine | 303.1187 | Asparagine-D |
| Asparagine | 257.0769 | Asparagine-NL |
| Asparagine | 473.1766 | Asparagine-D-D |
| Asparagine | 427.1348 | Asparagine-D-NL |
| Asparagine | 301.1041 | Asparagine-D-neg |
| Asparagine | 133.0608 | Asparagine-I-pos |
| Asparagine | 131.0462 | Asparagine-I-neg |
| Aspartic acid | 304.1027 | Aspartic acid-D |
| Aspartic acid | 258.0609 | Aspartic acid-NL |
| Aspartic acid | 302.0881 | Aspartic acid-D-neg |
| Aspartic acid | 134.0448 | Aspartic acid-I-pos |
| Aspartic acid | 132.0302 | Aspartic acid-I-neg |
| Betaine | 118.0863 | Betaine-I-pos |
| Caffeine | 365.1456 | Caffeine-D |
| Caffeine | 195.0877 | Caffeine-I-pos |
| Choline | 104.107 | Choline-I-pos |
| Citrulline | 346.1609 | Citrulline-D |
| Citrulline | 300.1191 | Citrulline-NL |
| Citrulline | 516.2188 | Citrulline-D-D |
| Citrulline | 472.1926 | Citrulline-D-NL |

| Compound | m/z values | Ion name |
|---|---|---|
| Citrulline | 344.1463 | Citrulline-D-neg |
| Citrulline | 176.103 | Citrulline-I-pos |
| Citrulline | 174.0884 | Citrulline-I-neg |
| Cysteine | 292.085 | Cysteine-D |
| Cysteine | 246.0431 | Cysteine-NL |
| Cysteine | 462.1429 | Cysteine-D-D |
| Cysteine | 416.101 | Cysteine-D-NL |
| Cysteine | 290.0703 | Cysteine-D-neg |
| Cysteine | 122.0271 | Cysteine-I-pos |
| Cysteine | 120.0124 | Cysteine-I-neg |
| DOPA | 368.134 | DOPA-D |
| DOPA | 322.0922 | DOPA-NL |
| DOPA | 366.1194 | DOPA-D-neg |
| DOPA | 198.0761 | DOPA-I-pos |
| DOPA | 196.0615 | DOPA-I-neg |
| Dopamine | 324.1442 | Dopamine-D |
| Dopamine | 278.1023 | Dopamine-NL |
| Dopamine | 322.1296 | Dopamine-D-neg |
| Dopamine | 154.0863 | Dopamine-I-pos |
| Dopamine | 152.0717 | Dopamine-I-neg |
| GABA | 274.1286 | GABA-D |
| GABA | 228.0867 | GABA-NL |
| GABA | 272.1139 | GABA-D-neg |
| GABA | 104.0707 | GABA-I-pos |
| GABA | 102.056 | GABA-I-neg |
| Glutamic acid | 318.1184 | Glutamic acid-D |
| Glutamic acid | 272.0765 | Glutamic acid-NL |
| Glutamic acid | 316.1037 | Glutamic acid-D-neg |
| Glutamic acid | 148.0605 | Glutamic acid-I-pos |
| Glutamic acid | 146.0458 | Glutamic acid-I-neg |
| Glutamine | 317.1344 | Glutamine-D |

| Compound | m/z values | Ion name |
|---|---|---|
| Glutamine | 271.0925 | Glutamine-NL |
| Glutamine | 315.1197 | Glutamine-D-neg |
| Glutamine | 147.0765 | Glutamine-I-pos |
| Glutamine | 145.0618 | Glutamine-I-neg |
| Glycine | 246.0973 | Glycine-D |
| Glycine | 200.0554 | Glycine-NL |
| Glycine | 244.0826 | Glycine-D-neg |
| Glycine | 76.0394 | Glycine-I-pos |
| Glycine | 74.0247 | Glycine-I-neg |
| Histamine | 282.1449 | Histamine-D |
| Histamine | 236.103 | Histamine-NL |
| Histamine | 280.1302 | Histamine-D-neg |
| Histamine | 112.087 | Histamine-I-pos |
| Histamine | 110.0723 | Histamine-I-neg |
| Histidine | 326.1347 | Histidine-D |
| Histidine | 280.0928 | Histidine-NL |
| Histidine | 496.1926 | Histidine-D-D |
| Histidine | 450.1508 | Histidine-D-NL |
| Histidine | 324.1201 | Histidine-D-neg |
| Histidine | 156.0768 | Histidine-I-pos |
| Histidine | 154.0622 | Histidine-I-neg |
| Homocitrulline | 360.1766 | Homocitrulline-D |
| Homocitrulline | 314.1347 | Homocitrulline-NL |
| Homocitrulline | 530.2345 | Homocitrulline-D-D |
| Homocitrulline | 484.1926 | Homocitrulline-D-NL |
| Homocitrulline | 358.1619 | Homocitrulline-D-neg |
| Homocitrulline | 190.1187 | Homocitrulline-I-pos |
| Homocitrulline | 188.104 | Homocitrulline-I-neg |
| Hypoxanthine | 307.1037 | Hypoxanthine-D |
| Hypoxanthine | 263.0775 | Hypoxanthine-NL |
| Hypoxanthine | 305.0891 | Hypoxanthine-D-neg |

| Compound | m/z values | Ion name |
|---|---|---|
| Hypoxanthine | 137.0458 | Hypoxanthine-I-pos |
| Hypoxanthine | 135.0312 | Hypoxanthine-I-neg |
| Kynurenine | 379.15 | Kynurenine-D |
| Kynurenine | 333.1082 | Kynurenine-NL |
| Kynurenine | 549.2079 | Kynurenine-D-D |
| Kynurenine | 503.1661 | Kynurenine-D-NL |
| Kynurenine | 377.1354 | Kynurenine-D-neg |
| Kynurenine | 547.1933 | Kynurenine-D-D-neg |
| Kynurenine | 209.0921 | Kynurenine-I-pos |
| Kynurenine | 207.0775 | Kynurenine-I-neg |
| (Iso)Leucine | 302.1599 | (Iso)Leucine-D |
| (Iso)Leucine | 258.1336 | (Iso)Leucine-NL |
| (Iso)Leucine | 300.1452 | (Iso)Leucine-D-neg |
| (Iso)Leucine | 132.102 | (Iso)Leucine-I-pos |
| (Iso)Leucine | 130.0873 | (Iso)Leucine-I-neg |
| Lysine | 317.1708 | Lysine-D |
| Lysine | 271.1289 | Lysine-NL |
| Lysine | 487.2287 | Lysine-D-D |
| Lysine | 441.1868 | Lysine-D-NL |
| Lysine | 315.1561 | Lysine-D-neg |
| Lysine | 485.214 | Lysine-D-D-neg |
| Lysine | 147.1129 | Lysine-I-pos |
| Lysine | 145.0982 | Lysine-I-neg |
| Methionine | 320.1163 | Methionine-D |
| Methionine | 274.0744 | Methionine-NL |
| Methionine | 318.1016 | Methionine-D-neg |
| Methionine | 150.0584 | Methionine-I-pos |
| Methionine | 148.0437 | Methionine-I-neg |
| NH3 | 188.0918 | NH3-D |
| NH3 | 142.0499 | NH3-NL |
| NH3 | 358.1497 | NH3-D-D |

| Compound | m/z values | Ion name |
|---|---|---|
| NH3 | 312.1078 | NH3-D-NL |
| NH3 | 186.0771 | NH3-D-neg |
| Niacinamide | 293.1132 | Niacinamide-D |
| Niacinamide | 247.0714 | Niacinamide-NL |
| Niacinamide | 123.0553 | Niacinamide-I-pos |
| Ornithine | 303.1551 | Ornithine-D |
| Ornithine | 257.1132 | Ornithine-NL |
| Ornithine | 473.213 | Ornithine-D-D |
| Ornithine | 427.1712 | Ornithine-D-NL |
| Ornithine | 301.1405 | Ornithine-D-neg |
| Ornithine | 471.1984 | Ornithine-D-D-neg |
| Ornithine | 133.0972 | Ornithine-I-pos |
| Ornithine | 131.0826 | Ornithine-I-neg |
| Orotic acid | 327.0823 | Orotic acid-D |
| Orotic acid | 497.1402 | Orotic acid-D-D |
| Orotic acid | 325.0677 | Orotic acid-D-neg |
| Orotic acid | 157.0244 | Orotic acid-I-pos |
| Orotic acid | 155.0098 | Orotic acid-I-neg |
| Phenylalanine | 336.1442 | Phenylalanine-D |
| Phenylalanine | 290.1023 | Phenylalanine-NL |
| Phenylalanine | 334.1296 | Phenylalanine-D-neg |
| Phenylalanine | 166.0863 | Phenylalanine-I-pos |
| Phenylalanine | 164.0717 | Phenylalanine-I-neg |
| Proline | 286.1286 | Proline-D |
| Proline | 242.1023 | Proline-NL |
| Proline | 284.1139 | Proline-D-neg |
| Proline | 116.0707 | Proline-I-pos |
| Proline | 114.056 | Proline-I-neg |
| Putrescine | 259.1653 | Putrescine-D |
| Putrescine | 213.1234 | Putrescine-NL |
| Putrescine | 429.2232 | Putrescine-D-D |

| Compound | m/z values | Ion name |
|---|---|---|
| Putrescine | 383.1813 | Putrescine-D-NL |
| Putrescine | 89.1074 | Putrescine-I-pos |
| Putrescine | 87.0927 | Putrescine-I-neg |
| Riboflavin | 547.2035 | Riboflavin-D |
| Riboflavin | 545.1889 | Riboflavin-D-neg |
| Riboflavin | 377.1456 | Riboflavin-I-pos |
| Riboflavin | 375.131 | Riboflavin-I-neg |
| SAM | 569.2025 | SAM-D |
| SAM | 739.2604 | SAM-D-D |
| SAM | 399.1446 | SAM-I-pos |
| SAM | 285.1049 | SAM-D-D-2+ |
| SAM | 398.1372 | SAM-I-neg |
| Serine | 276.1078 | Serine-D |
| Serine | 230.066 | Serine-NL |
| Serine | 274.0932 | Serine-D-neg |
| Serine | 106.0499 | Serine-I-pos |
| Serine | 104.0353 | Serine-I-neg |
| Serotonin | 347.1602 | Serotonin-D |
| Serotonin | 301.1183 | Serotonin-NL |
| Serotonin | 345.1455 | Serotonin-D-neg |
| Serotonin | 177.1023 | Serotonin-I-pos |
| Serotonin | 175.0876 | Serotonin-I-neg |
| Spermidine | 316.2231 | Spermidine-D |
| Spermidine | 270.1813 | Spermidine-NL |
| Spermidine | 486.281 | Spermidine-D-D |
| Spermidine | 440.2392 | Spermidine-D-NL |
| Spermidine | 314.2085 | Spermidine-D-neg |
| Spermidine | 146.1652 | Spermidine-I-pos |
| Spermidine | 144.1506 | Spermidine-I-neg |
| Spermine | 373.281 | Spermine-D |
| Spermine | 327.2391 | Spermine-NL |

| Compound | m/z values | Ion name |
|---|---|---|
| Spermine | 543.3389 | Spermine-D-D |
| Spermine | 497.297 | Spermine-D-NL |
| Spermine | 371.2663 | Spermine-D-neg |
| Spermine | 203.2231 | Spermine-I-pos |
| Spermine | 201.2084 | Spermine-I-neg |
| TMAO | 230.1387 | TMAO-D |
| TMAO | 115.573 | TMAO-NL |
| TMAO | 76.0757 | TMAO-I-pos |
| Taurine | 296.0799 | Taurine-D |
| Taurine | 250.038 | Taurine-NL |
| Taurine | 294.0652 | Taurine-D-neg |
| Taurine | | Taurine-I-pos |
| Taurine | 124.0073 | Taurine-I-neg |
| Threonine | 290.1235 | Threonine-D |
| Threonine | 244.0816 | Threonine-D-NL |
| Threonine | 288.1088 | Threonine-D-neg |
| Threonine | 120.0656 | Threonine-I-pos |
| Threonine | 118.0509 | Threonine-I-neg |
| Tryptamine | 331.1653 | Tryptamine-D |
| Tryptamine | 285.1234 | Tryptamine-NL |
| Tryptamine | 501.2232 | Tryptamine-D-D |
| Tryptamine | 455.1813 | Tryptamine-D-NL |
| Tryptamine | 329.1506 | Tryptamine-D-neg |
| Tryptamine | 161.1074 | Tryptamine-I-pos |
| Tryptamine | 159.0927 | Tryptamine-I-neg |
| Tryptophan | | Tryptophan-D |
| Tryptophan | 329.1132 | Tryptophan-NL |
| Tryptophan | 373.1405 | Tryptophan-D-neg |
| Tryptophan | 205.0972 | Tryptophan-I-pos |
| Tryptophan | 203.0826 | Tryptophan-I-neg |
| Tyrosine | 352.1391 | Tyrosine-D |

| Compound | m/z values | Ion name |
|---|---|---|
| Tyrosine | 306.0973 | Tyrosine-NL |
| Tyrosine | 350.1245 | Tyrosine-D-neg |
| Tyrosine | 182.0812 | Tyrosine-I-pos |
| Tyrosine | 180.0666 | Tyrosine-I-neg |
| Uracil | 283.0925 | Uracil-D |
| Uracil | 453.1504 | Uracil-D-D |
| Uracil | 113.0346 | Uracil-I-pos |
| Uridine | 415.1348 | Uridine-D |
| Uridine | 413.1201 | Uridine-D-neg |
| Uridine | 371.1086 | Uridine-NL |
| Uridine | 245.0769 | Uridine-I-pos |
| Uridine | 243.0622 | Uridine-I-neg |
| Urea | 231.0976 | Urea-D |
| Urea | 185.0557 | Urea-NL |
| Urea | 401.1555 | Urea-D-D |
| Urea | 355.1136 | Urea-D-NL |
| Urea | 229.0829 | Urea-D-neg |
| Urea | 61.0397 | Urea-I-pos |
| Urea | 59.025 | Urea-I-neg |
| Valine | 288.1442 | Valine-D |
| Valine | 242.1023 | Valine-NL |
| Valine | 286.1296 | Valine-D-neg |
| Valine | 118. | Valine-I-pos |
| Valine | 116.0177 | Valine-I-neg |
| Carbamoyl phosphate | 312.0479 | Carbamoyl phosphate-D |
| Carbamoyl phosphate | 266.0061 | Carbamoyl phosphate-NL |
| Carbamoyl phosphate | 310.0333 | Carbamoyl phosphate-D-neg |
| Carbamoyl phosphate | 141.99 | Carbamoyl phosphate-I-pos |
| Carbamoyl phosphate | 139.9754 | Carbamoyl phosphate-I-neg |
| Aminoadipic acid | 332.134 | Aminoadipic acid-D |
| Aminoadipic acid | 286.0922 | Aminoadipic acid-NL |

| Compound | m/z values | Ion name |
|---|---|---|
| Aminoadipic acid | 330.1194 | Aminoadipic acid-D-neg |
| Aminoadipic acid | 162.0761 | Aminoadipic acid-I-pos |

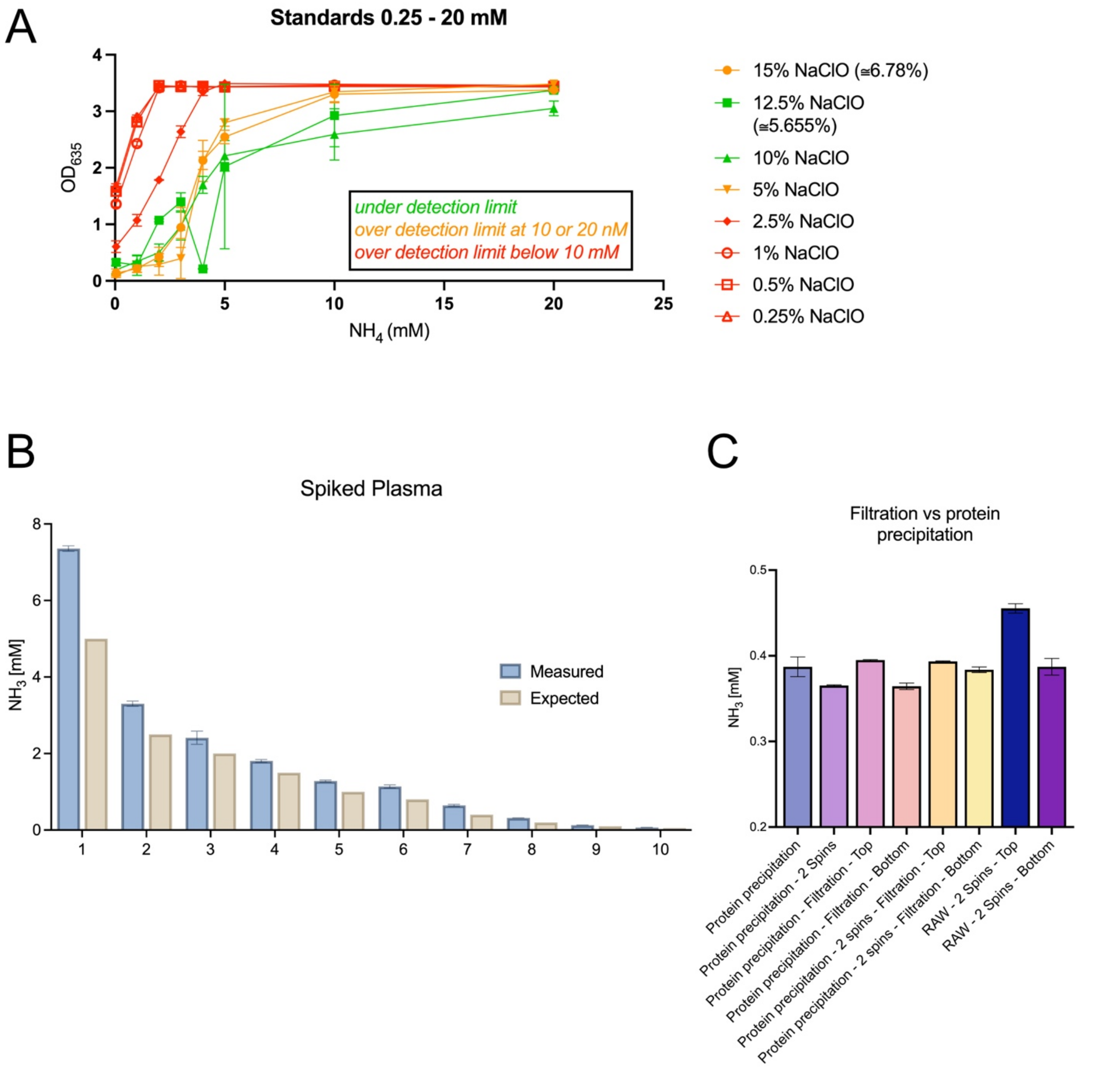


**Fig. S1. Evaluation of colorimetric ammonia quantification assays and sample preparation effects.** (A) Calibration curves obtained using ammonium chloride standards (0.25–20 mM) prepared with varying sodium hypochlorite (NaClO) concentrations. Increasing oxidant concentration reduced the dynamic range, with higher NaClO (>5%) leading to signal saturation or loss of linearity. Lines and error bars represent mean ± SD of technical replicates. (B) Measured versus expected ammonia concentrations in plasma samples spiked with known amounts of ammonium chloride, showing systematic overestimation of concentrations by the colorimetric assay despite protein precipitation. Bars indicate mean ± SD. (C) Comparison of ammonia quantification following different sample-preparation procedures, including protein precipitation, filtration, and combinations thereof. “RAW” indicates untreated plasma. Filtration and protein precipitation yield comparable results, with minimal variation between top and bottom fractions after centrifugation.

A

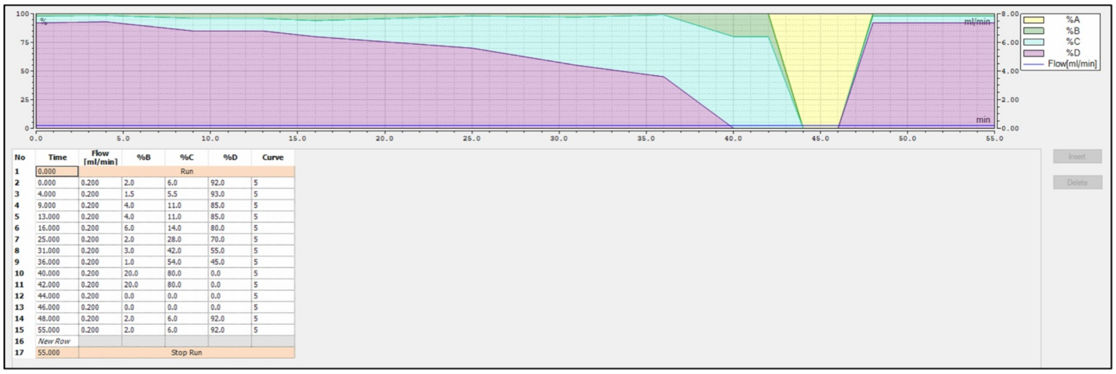


| No | Time | Flow [ml/min] | %B | %C | %D | Curve |
|---|---|---|---|---|---|---|
| 1 | 0.000 | Run | | | | |
| 2 | 0.000 | 0.200 | 2.0 | 6.0 | 92.0 | 5 |
| 3 | 4.000 | 0.200 | 1.5 | 5.5 | 93.0 | 5 |
| 4 | 9.000 | 0.200 | 4.0 | 11.0 | 85.0 | 5 |
| 5 | 13.000 | 0.200 | 4.0 | 11.0 | 85.0 | 5 |
| 6 | 16.000 | 0.200 | 6.0 | 14.0 | 80.0 | 5 |
| 7 | 25.000 | 0.200 | 2.0 | 28.0 | 70.0 | 5 |
| 8 | 31.000 | 0.200 | 3.0 | 42.0 | 55.0 | 5 |
| 9 | 36.000 | 0.200 | 1.0 | 54.0 | 45.0 | 5 |
| 10 | 40.000 | 0.200 | 20.0 | 80.0 | 0.0 | 5 |
| 11 | 42.000 | 0.200 | 20.0 | 80.0 | 0.0 | 5 |
| 12 | 44.000 | 0.200 | 0.0 | 0.0 | 0.0 | 5 |
| 13 | 46.000 | 0.200 | 0.0 | 0.0 | 0.0 | 5 |
| 14 | 48.000 | 0.200 | 2.0 | 6.0 | 92.0 | 5 |
| 15 | 55.000 | 0.200 | 2.0 | 6.0 | 92.0 | 5 |
| 16 | New Row | | | | | |
| 17 | 55.000 | Stop Run | | | | |

B

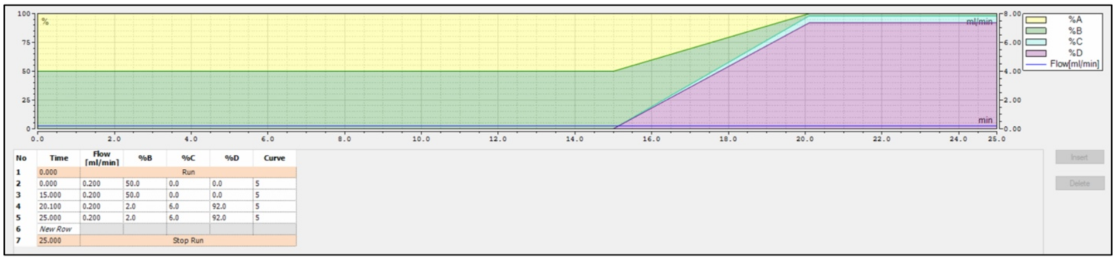


| No | Time | Flow [ml/min] | %B | %C | %D | Curve |
|---|---|---|---|---|---|---|
| 1 | 0.000 | Run | | | | |
| 2 | 0.000 | 0.200 | 50.0 | 0.0 | 0.0 | 5 |
| 3 | 15.000 | 0.200 | 50.0 | 0.0 | 0.0 | 5 |
| 4 | 20.100 | 0.200 | 2.0 | 6.0 | 92.0 | 5 |
| 5 | 25.000 | 0.200 | 2.0 | 6.0 | 92.0 | 5 |
| 6 | New Row | | | | | |
| 7 | 25.000 | Stop Run | | | | |

C

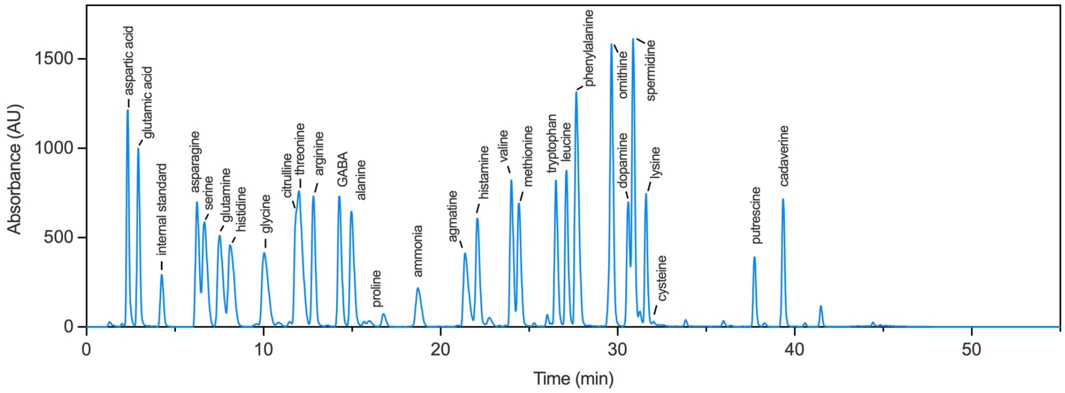


**Fig. S2. LC gradient optimization and chromatographic separation of DEEMM-derivatized nitrogen metabolites.** (A) Elution gradient used for metabolite separation. (B) Washing gradient used to clean and re-equilibrate the column between runs. Eluents: A: dd$H_2O$; B: 99.9% HPLC-grade methanol; C: 99.9% HPLC-grade acetonitrile; D: freshly prepared, sterile-filtered acetate buffer. (C) Labeled LC–UV chromatogram showing the elution profile of the standard mix used for quantification of DEEMM aminoenone derivatives of nitrogen-containing metabolit, separated over a 55-min run.

**Table S2. Elution gradient used for LC separation of DEEMM-derivatized nitrogen metabolites.** Gradient program detailing the composition of eluents B–D over time, used for the chromatographic separation shown in Figure S2. The flow rate was maintained at 0.2 mL min$^{-1}$ throughout the run. Eluents were: (A) dd$H_2O$, (B) 99.9% HPLC-grade methanol, (C) 99.9% HPLC-grade acetonitrile, and (D) freshly prepared, sterile-filtered acetate buffer.

| Time [min] | Flow [ml/min] | %B | %C | %D | Curve |
|---|---|---|---|---|---|
| 0.0 | 0.200 | 2.0 | 6.0 | 92.0 | 5 |
| 4.0 | 0.200 | 1.5 | 5.5 | 93.0 | 5 |
| 9.0 | 0.200 | 4.0 | 11.0 | 85.0 | 5 |
| 13.0 | 0.200 | 4.0 | 11.0 | 85.0 | 5 |
| 16.0 | 0.200 | 6.0 | 14.0 | 80.0 | 5 |
| 25.0 | 0.200 | 2.0 | 28.0 | 70.0 | 5 |
| 31.0 | 0.200 | 3.0 | 42.0 | 55.0 | 5 |
| 36.0 | 0.200 | 1.0 | 54.0 | 45.0 | 5 |
| 40.0 | 0.200 | 20.0 | 80.0 | 0.0 | 5 |
| 42.0 | 0.200 | 20.0 | 80.0 | 0.0 | 5 |
| 44.0 | 0.200 | 0.0 | 0.0 | 0.0 | 5 |
| 46.0 | 0.200 | 0.0 | 0.0 | 0.0 | 5 |
| 48.0 | 0.200 | 2.0 | 6.0 | 92.0 | 5 |
| 55.0 | 0.200 | 2.0 | 6.0 | 92.0 | 5 |

**Table S3. Washing gradient used to clean and re-equilibrate the column between analytical runs.** The flow rate was maintained at 0.2 mL min$^{-1}$. Eluents were: (A) dd$H_2O$, (B) 99.9% HPLC-grade methanol, (C) 99.9% HPLC-grade acetonitrile, and (D) freshly prepared, sterile-filtered acetate buffer. “Curve 5” corresponds to a linear gradient transition.

| Time [min] | Flow [ml/min] | %B | %C | %D | Curve |
|---|---|---|---|---|---|
| 0.0 | 0.200 | 50.0 | 0.0 | 0.0 | 5 |

| Time [min] | Flow [ml/min] | %B | %C | %D | Curve |
|---|---|---|---|---|---|
| 4.0 | 0.200 | 50.0 | 0.0 | 0.0 | 5 |
| 9.0 | 0.200 | 2.0 | 6.0 | 92.0 | 5 |
| 13.0 | 0.200 | 2.0 | 6.0 | 92.0 | 5 |

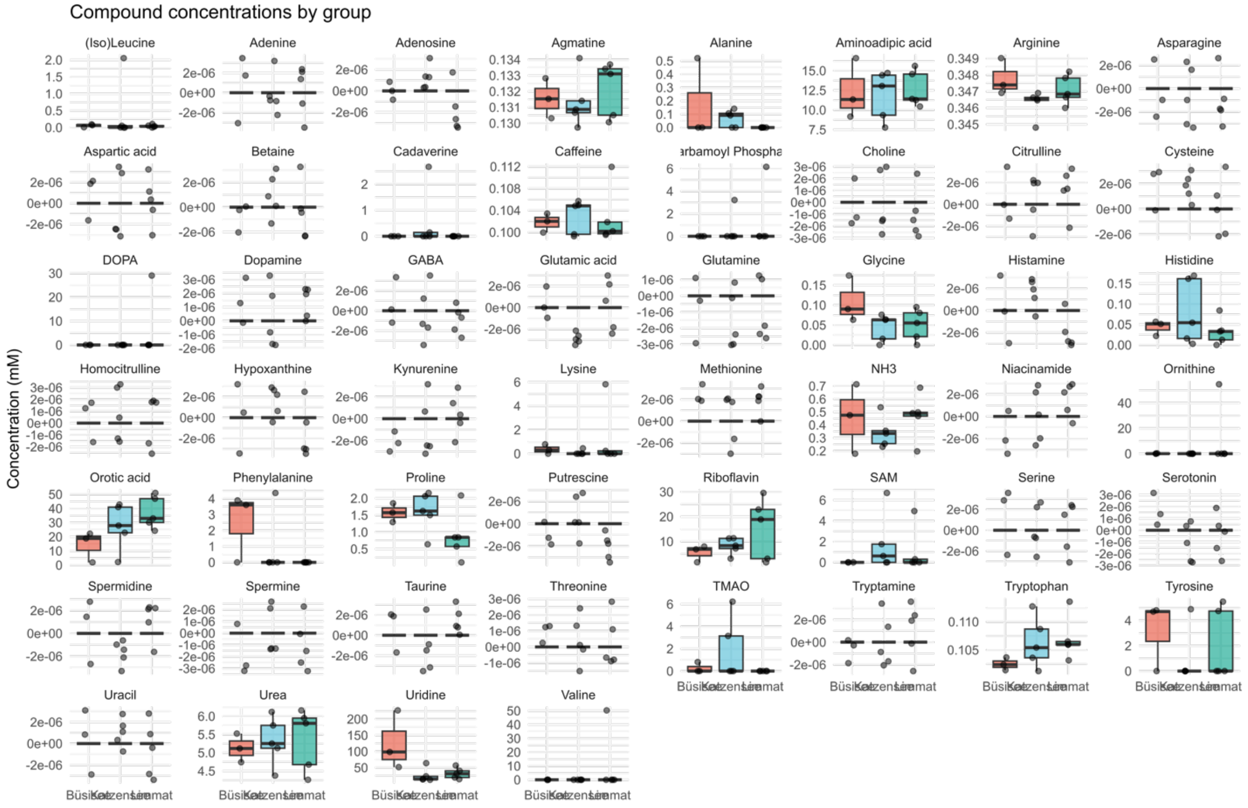


**Figure S3. Quantification of nitrogen-containing metabolites in freshwater samples.**

Boxplots showing the concentrations of all compounds quantified using the LC–UV method described in this study across the different freshwater sample groups. Each point represents an independent sample, and boxes indicate the interquartile range with median lines.
